# Evolution of the gene regulatory network of body axis by enhancer hijacking in amphioxus

**DOI:** 10.1101/2023.06.11.544506

**Authors:** Chenggang Shi, Shuang Chen, Huimin Liu, Shiqi Li, Yanhui Wang, Rongrong Pan, Xiaotong Wu, Xuewen Li, Jingjing Li, Chaofan Xing, Xian Liu, Yiquan Wang, Qingming Qu, Guang Li

## Abstract

A central goal of evolutionary developmental biology is to decipher the evolutionary pattern of gene regulatory networks (GRNs) that control morphogenesis during embryonic development, and the underlying molecular mechanism of the GRNs evolution. The Nodal signaling that governs the body axes of deuterostomes exhibits a conserved GRN orchestrated principally by Nodal, Gdf1/3 and Lefty. Despite that the GRN retains a conserved function in deuterostomes, the expression patterns of *Gdf1/3* and *Nodal* are derived in cephalochordate amphioxus, implying a specific rewiring of the GRN in this lineage. Here we examined the regulatory mechanism and evolution of this GRN in amphioxus by functional genetic manipulations. We found that while the amphioxus *Gdf1/3* orthologue shows nearly no expression during embryogenesis, its duplicate *Gdf1/3-like* linking to *Lefty* is zygotically expressed in a similar pattern as *Lefty*. Mutant and transgenic analyses revealed that *Gdf1/3* is no longer crucial for amphioxus axial development. Instead, *Gdf1/3-like* assumes this responsibility, likely through hijacking *Lefty* enhancers. We also showed that amphioxus Nodal has become an indispensable maternal factor to compensate for the loss of maternal *Gdf1/3* expression. We therefore demonstrated a case that the evolution of an ancestral GRN could be triggered by enhancer hijacking events. This pivotal event has allowed the emergence of a new GRN in the extant amphioxus, presumably evolving through a stepwise process. The co-expression of *Gdf1/3-like* and *Lefty* achieved by shared regulatory region may have provided developmental robustness during body axis formation in extant amphioxus, which provides a selection-based hypothesis for the phenomena called developmental system drift.

## Introduction

The developmental regulatory mechanisms of body axis formation have been investigated in numerous metazoans, which have greatly advanced evolutionary developmental biology (evo-devo) as a research field^1^. Importantly, these accumulated data also make it possible to analyze how the gene regulatory networks (GRNs) controlling body axis has been evolving in different clades, a key theme in evo-devo^1,2^. Nevertheless, detailed functional genetic evidence showing how GRNs could be rewired during evolution is usually lacking, which hinders our understanding of the evolvability of organisms.

Nodal signaling plays a conserved role in patterning the dorsal-ventral (D-V) and left-right (L-R) axis in deuterostomes, exhibiting a conserved GRN orchestrated principally by Nodal, Gdf1/3 and Lefty^3^. The expression patterns of genes coding Nodal and Gdf1/3 are highly conserved in echinoderms and vertebrates, with *Nodal* being expressed zygotically (unilaterally at neurula or larva stage) (Supplementary Table 1), and *Gdf1/3* both maternally and zygotically (but bilaterally at neurula stage) (Supplementary Table 2). Zygotic Nodal functions synergistically with preexisting maternal Gdf1/3 by forming heterodimers to activate the signaling pathway^4,5^. Robust Nodal signaling is safeguarded by a positive feedback loop from further activation of *Nodal* itself and a negative feedback loop through inducing the expression of *Lefty* encoding an inhibitor of the signaling ^6^.

Previous studies have identified one *Nodal,* one *Lefty* and two *Gdf1/3* (tentatively named *Gdf1/3-like1/Vg1* and *Gdf1/3-like2*) genes in basal chordate amphioxus^7-9^. The expression of *Nodal*, *Lefty* and one of *Gdf1/3* (known as *Gdf1/3-like1* or *Vg1*) genes have been investigated^8-12^. Curiously, it was found that *Nodal* and *Gdf1/3-like1* are both maternally supplied and zygotically expressed unilaterally at neurula stage (Supplementary Table 1 and 2). This implies that the regulatory network governing the Nodal signaling pathway has undergone alterations within this particular clade, while its function in D-V and L-R axes patterning has been preserved^9,10,13,14^. We thus speculate that the Nodal signaling of amphioxus would be an excellent case to trace the evolutionary history of a GRN and to clarify the mechanism underlying it. To evaluate this, we analyzed the GRN of amphioxus Nodal signaling using mutant and transgenic lines and dissected its evolutionary history by integrating available functional genetic data from other deuterostomes.

## Results

### Evolution of the two *Gdf1/3* genes in amphioxus

*Gdf1/3* genes have only been detected in deuterostomes^7,15-18^. The gene is linked to *Bmp2/4* in many deuterostome species (Fig. 1a), and also shows the most sequence similarity to *Bmp2/4*^7,19^. Accordingly, *Gdf1/3* is thought to have originated from *Bmp2/4* by a tandem duplication event that occurred in the common ancestor of deuterostomes (Fig. 1a)^18,19^. However, Floridae amphioxus (*Branchiostoma floridae*) was reported to contain two *Gdf1/3* genes^7^, one of which (previously named *GDF1/3-like2* by Satou et al^7^) is linked to *Bmp2/4* as was found in other deuterostomes, representing the ancestral amphioxus *Gdf1/3* gene (hereafter renamed *Gdf1/3*, Fig. 1a). Intriguingly, the other one (previously known as *GDF1/3-like1*^7^ or *Vg1*^9^, here renamed as *Gdf1/3-like*, Fig. 1a) is also linked to a TGF-β family gene, *Lefty*^7^. This unusual gene arrangement, together with the view that *Lefty* originated in deuterostomes at that time, led Satou and his colleagues proposing that the *Gdf1/3-like*-*Lefty* gene pair derived from either a duplication of the *Gdf1/3*-*Bmp2/4* gene pair or two duplications of *Gdf1/3* (*Gdf1/3* duplicated first to generate *Gdf1/3-like*, which then translocated and duplicated tandemly to generate *Lefty*)^7^. However, this view was not supported by molecular phylogenetic analysis^7^ and was rejected by the recent finding of *Lefty* genes in Lophotrochozoa^20^. Further survey on recently updated genomes of bilaterians, especially those reported to have *Lefty* genes, revealed that the *Gdf1/3-like* gene and its linkage to *Lefty* exist only in amphioxus species, but not in any other bilaterians examined (Fig. 1a). These findings suggest that the *Gdf1/3-like* gene most likely arose in Cephalochordata, or at least in the genus of *Brachiostoma*, through a tandem duplication of *Gdf1/3*, followed by a translocation of it to the *Lefty* locus (Fig. 1a). In line with this proposal, lineage-specific duplication of the *Gdf1/3* gene and translocation of the duplicate to other genomic regions have also been found in at least two lineages of vertebrates^17^.

**Figure 1.**
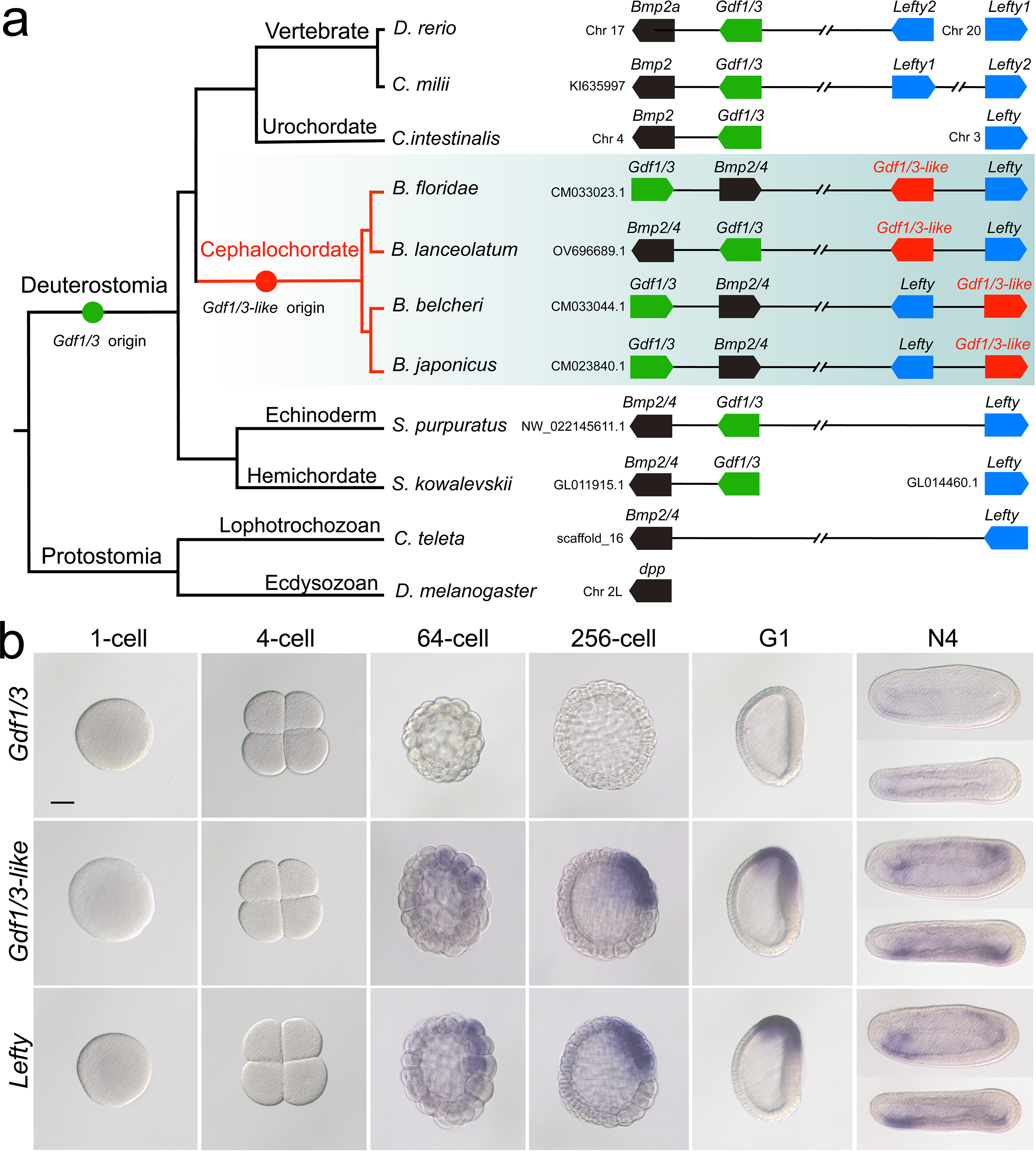
Synteny and expression pattern of amphioxus *Gdf1/3* and *Gdf1/3-like* gene. **(a)** Arrangement of *Gdf1/3*, *Bmp2/4*, *Gdf1/3-like* and *Lefty* genes in representative bilaterian genomes. Black lines represent the genes (boxes) at both ends are tightly linked and fractured lines represent the genes are close together on a same chromosome or scafford indicated. zebrafish *(D. rerio*), elephant Shark (*C. milii*), vase tunicate (*Ciona intestinalis*), Florida amphioxus (*B. floridae*), European amphioxus (*B. lanceolatum*), Asia amphioxus (*B. belcheri* and *B. japonicum*), sea urchin (*S. purpuratus*), acorn worm (*Saccoglossus kowalevskii*), polychaete worm (*Capitella teleta*), fruit fly (*Drosophila melanogaster*). **(b)** The spatio-temporal expression pattern of *Gdf1/3*, *Gdf1/3-like* and *Lefty* at different stages of *B. floridae* embryos. Embryos at 64-cell to G1 stage were viewed from the left side with anterior to the left, and those at N4 stage were viewed from either the left site (upper panels) or dorsal side (lower panels) with anterior to the left. Scale bar, 50 μm.

### Amphioxus *Gdf1/3* lost the ancestral role in body axes formation

We next asked if the *Gdf1/3* in amphioxus is functionally similar to its orthologue in vertebrates and sea urchins. We first analyzed its embryonic expression pattern using both *in situ* hybridization and qPCR. Unexpectedly, no *Gdf1/3* transcripts were detected in amphioxus embryos before the neurula stage, and very weak expression of the gene was found in few cells of the anterior ventral pharyngeal region during the late neurula and larva stages (Fig. 1b, Supplementary Fig. 1). We further generated amphioxus mutants of the *Gdf1/3* gene. Consistent with its restricted and weak expression pattern, the homozygous *Gdf1/3* mutants (*Gdf1/3^−/−^*) displayed normal D-V and L-R axis patterning as the wild type (WT) by the 3-gill slit stage (Supplementary Fig. 2). These data suggested that *Gdf1/3* has disassociated from the GRN of body axis formation in living amphioxus. It is, however, notable that overexpression of *Gdf1/3* by injecting its mRNA could result in the expansion of anterior and dorsal identity (expressing *FoxQ2*, *Wnt3*, *Brachyury* and *Chordin*) at the expense of ventral structures (expressing *Evx*) (Supplementary Fig. 3), as overactivation of the Nodal signaling^9,14^. Moreover, treatment with SB-505124, a selective inhibitor of Alk4/5/7, could ventralize the embryos injected (Supplementary Fig. 3). These results demonstrated that the Gdf1/3 still works as a ligand of Nodal signaling, although it is dispensable to the body axis formation of extant amphioxus.

### *Gdf1/3-like* is indispensable for body axes formation in amphioxus

As *Gdf1/3* is no longer involved in the body axis formation in amphioxus, while injection of *Gdf1/3* and *Gdf1/3-like* mRNA yielded similar phenotypes (Supplementary Fig. 3), we hypothesized that *Gdf1/3-like* is involved in the body axes formation. To test this, we first reanalyzed the expression pattern of *Gdf1/3-like* in amphioxus using different methods. Inconsistent with previous study^9^, no maternal expression of *Gdf1/3-like* was detected in our analyses (Fig. 1b, Supplementary Fig. 1). Zygotic expression of the gene was first detected in the dorsal blastomeres of the vegetal pole at the 32-64 cell stage, and then restricted in the dorsal blastoporal lip (Spemann organizer) at the gastrula stage and the left side of the embryo from the early neurula stage (Fig. 1b, Supplementary Fig. 1a). Notably, the expression pattern of *Gdf1/3-like* is similar to that of *Lefty* (Fig. 1b, Supplementary Fig. 1a).

We then created *Gdf1/3-like* mutants to see if it is required for amphioxus body axis development. *Gdf1/3-like^-/-^* embryos showed severe defects in axis formation. At the G5 stage, the *Gdf1/3-like^-/-^* embryos failed to flatten dorsally like the WT embryos (Supplementary Fig. 4a), and at subsequent stages they lacked most of the anterior and dorsal structures (Fig. 2a, Supplementary Fig. 4a). In addition, by the L2 stage, there were a minority of *Gdf1/3-like^+/-^* larvae showing no left but two-right phenotype (Fig. 2a). We further analyzed the expression patterns of the marker genes for various structures in the mutants. At the gastrulae (G5) stage of *Gdf1/3-like^-/-^*, the expression of *Gsc* in dorsal mesoderm, *Chordin* and *Netrin* in dorsal mesoderm and neural ectoderm and *SoxB1a* in dorsal pan-neural ectoderm disappeared, while that of *Evx* in ventral domain and *Ap2* in epidermal ectoderm expanded dorsally (Fig. 2b). At the T0 stage of *Gdf1/3-like^-/-^*, the expression of *Brachyury* in notochord, *FoxQ2* in anterior ectoderm, *Hex* in anterior endoderm, *m-actin* in somites, *Otx* in forebrain and anterior pharyngeal endoderm, and *Wnt3* in hindbrain and spinal cord were absent (Supplementary Fig. 4b). Moreover, in a minority of *Gdf1/3-like^+/-^* embryos of the T0-T1 stage, the left-sided expression of *Pit* in preoral pit and *Pou4* in oral primordium disappeared, while the right-sided expression of *Nkx2.1* in endostyle and *FoxE4* in club-shaped gland became bilaterally symmetrical (Fig. 2c). These data collectively indicated that *Gdf1/3-like* is indispensable for body axes formation in amphioxus and its loss-of-function leads to loss of dorsal, anterior and left identities.

**Figure 2.**
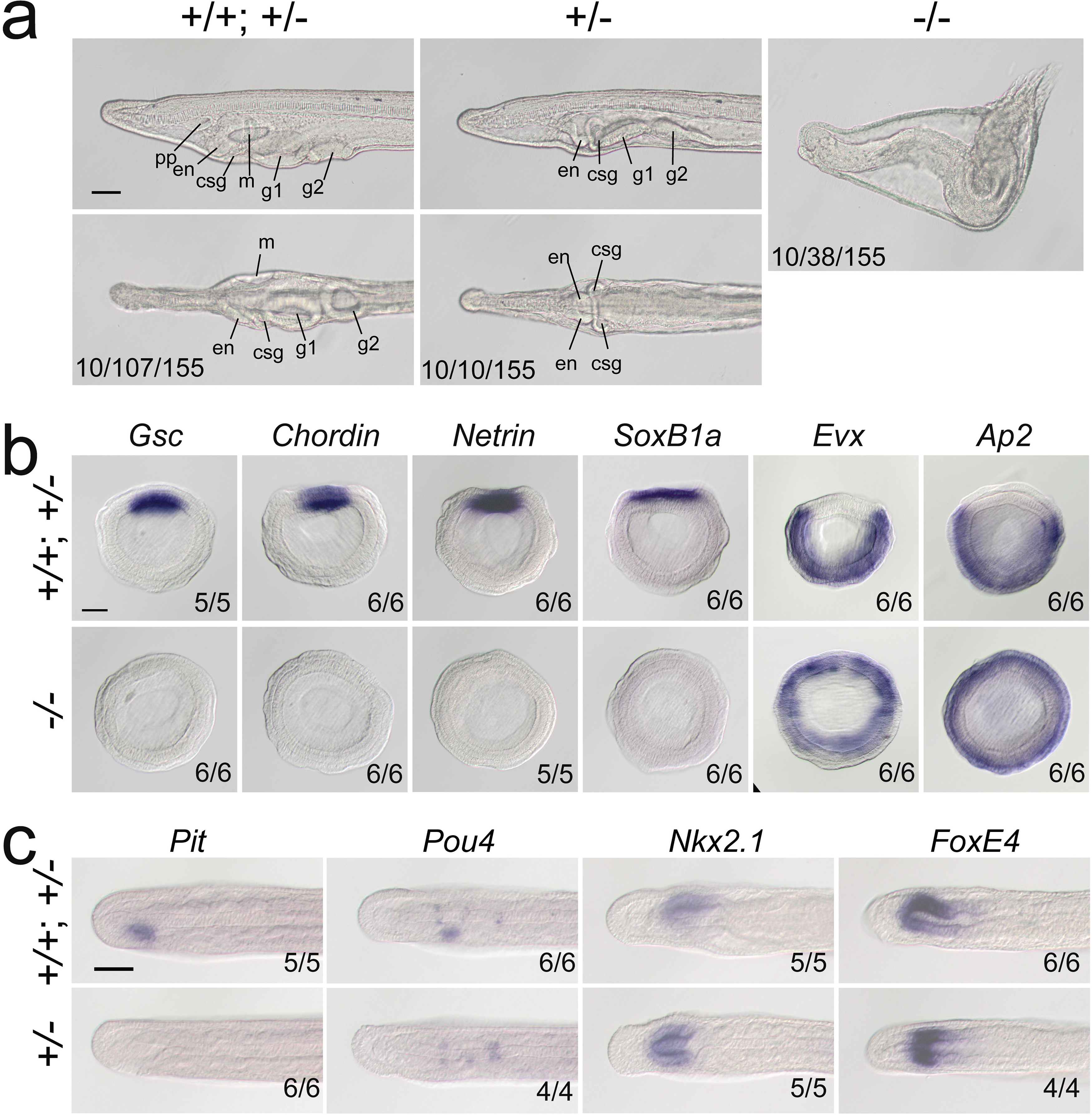
*Gdf1/3-like* loss-of-function affects amphioxus axes development. **(a)** Phenotypic analyses of *Gdf1/3-like* mutants. Larvae at L2 stage were observed from the left side (upper panels) and ventral side (under panels) with anterior to the left. pp, preoral pit; en, endostyle; csg, club shaped gland; m, mouth; g1, first gill slit; g2, second gill slit. The phenotypes of embryos at other stages are provided in Supplemental Figure 4. The three numbers (from left to right) at the bottom right of each panel indicate the number of larvae used for genotyping, the number of larvae with the phenotype, and the total number of larvae examined, respectively. **(b-c)** The expression of marker genes in *Gdf1/3-like* mutants. Embryos in (b) were at G5 stage and viewed from the blastopore with dorsal to the top, while embryos in (c) were at T0-T1 stage and viewed from the dorsal side with anterior to the left. Numbers at the bottom right indicate the number of times the genotype shown was identified, out of the total number of examined embryos or larvae with the expression pattern. Scale bars, 50 μm.

In vertebrates, transduction of Nodal signaling requires Gdf1/3 to form heterodimer with Nodal^4^. To see whether Gdf1/3-like is similarly required for Nodal signaling transduction in amphioxus, we examined the expression of *Nodal* and a target gene (*Gsc*) of the Nodal signaling in *Gdf1/3-like^-/-^* embryos. At the G1 stage, the expression of *Nodal* (most of them are probably maternal) was at a comparable level in *Gdf1/3-like^+/+^*, *Gdf1/3-like^+/^*^-^ and *Gdf1/3-like^-/-^* embryos (Supplementary Fig. 5). However, at the same stage the expression of *Gsc* was already activated in *Gdf1/3-like^+/+^* and most *Gdf1/3-like^+/^*^-^ but not in *Gdf1/3-like^-/-^* embryos (Supplementary Fig. 5). These results suggested that in the absence of Gdf1/3-like, Nodal alone could not activate the Nodal signaling in amphioxus.

### Maternal and zygotic Nodal is necessary for amphioxus body axes formation

In amphioxus, *Nodal* is expressed both maternally and zygotically^9^. To dissect its function during embryogenesis, we generated *Nodal* heterozygous animals and crossed them to analyze homozygous mutants. *Nodal^-/-^* embryos exhibited defects of L-R axis (Supplementary Fig. 6) as *Gdf1/3-like^+/-^*(shown above) and embryos in which late Nodal signaling activity was blocked^10,13^. This result showed that zygotic Nodal is necessary for L-R patterning in amphioxus.

However, these *Nodal*^-/-^ mutants do not show obvious dorsal-ventral or anterior-posterior defects (Supplementary Fig. 6). This is probably related to maternal *Nodal* expression. We therefore generated maternal *Nodal* mutants (M*Nodal*). Since zygotic *Nodal* mutants could not survive into adulthood, we screened for genetic mosaic females (founders) carrying oocytes of biallelic mutations^21^ at the *Nodal* locus (Supplementary Fig. 7a). Two such founders (named founder 1 and founder 2) were identified from nearly one hundred of animals. *In situ* hybridization experiment revealed that around 50% and 20% of eggs released by the founder 1 and 2 respectively showed no maternal *Nodal* mRNA accumulation (Fig. 3a-b). In line with this, when eggs of the two founders were fertilized with WT sperms (Supplementary Fig. 7b), around 50% and 20% of them (M*Nodal*) showed a mild ventralized phenotype respectively (Fig. 3c-d, Supplementary Fig. 7c). Among embryos generated from the above two crosses, approximately 50% and 20% of them respectively showed reduced expression of anterior and dorsal markers (*Gsc*, *Chordin*, *Netrin*, *SoxB1a*, *Brachyury*, *Wnt3*, *m-actin* and *FoxQ2*), and expanded expression of the ventral markers (*Evx* and *Ap2*) (Fig. 3e-f, Supplementary Fig. 8). We also crossed the two founders with male *Nodal^+/-^* animals to generate maternal and zygotic *Nodal* mutants (MZ*Nodal*). MZ*Nodal* embryos, as expected, accounting for around 25% and 10% of offspring of the founder 1 and 2 respectively, displayed ventralized phenotypes being more severe than M*Nodal* mutants, but comparable to *Gdf1/3* mutants (Fig. 3c-d, Supplementary Fig. 7c). Gene expression analysis showed that MZ*Nodal* mutants lost most of the dorsal and anterior identities, as indicated by loss of expressions of the dorsal and anterior marker genes (*Gsc*, *Chordin*, *Netrin*, *SoxB1a*, *Brachyury*, *Wnt3*, *m-actin* and *FoxQ2*), and expanded expressions of the ventral markers (*Evx* and *Ap2*) (Fig. 3e-f, Supplementary Fig. 8). These results collectively indicated that maternal *Nodal*, probably zygotic *Nodal* as well, is required for regulating the D-V axis formation in amphioxus.

**Figure 3.**
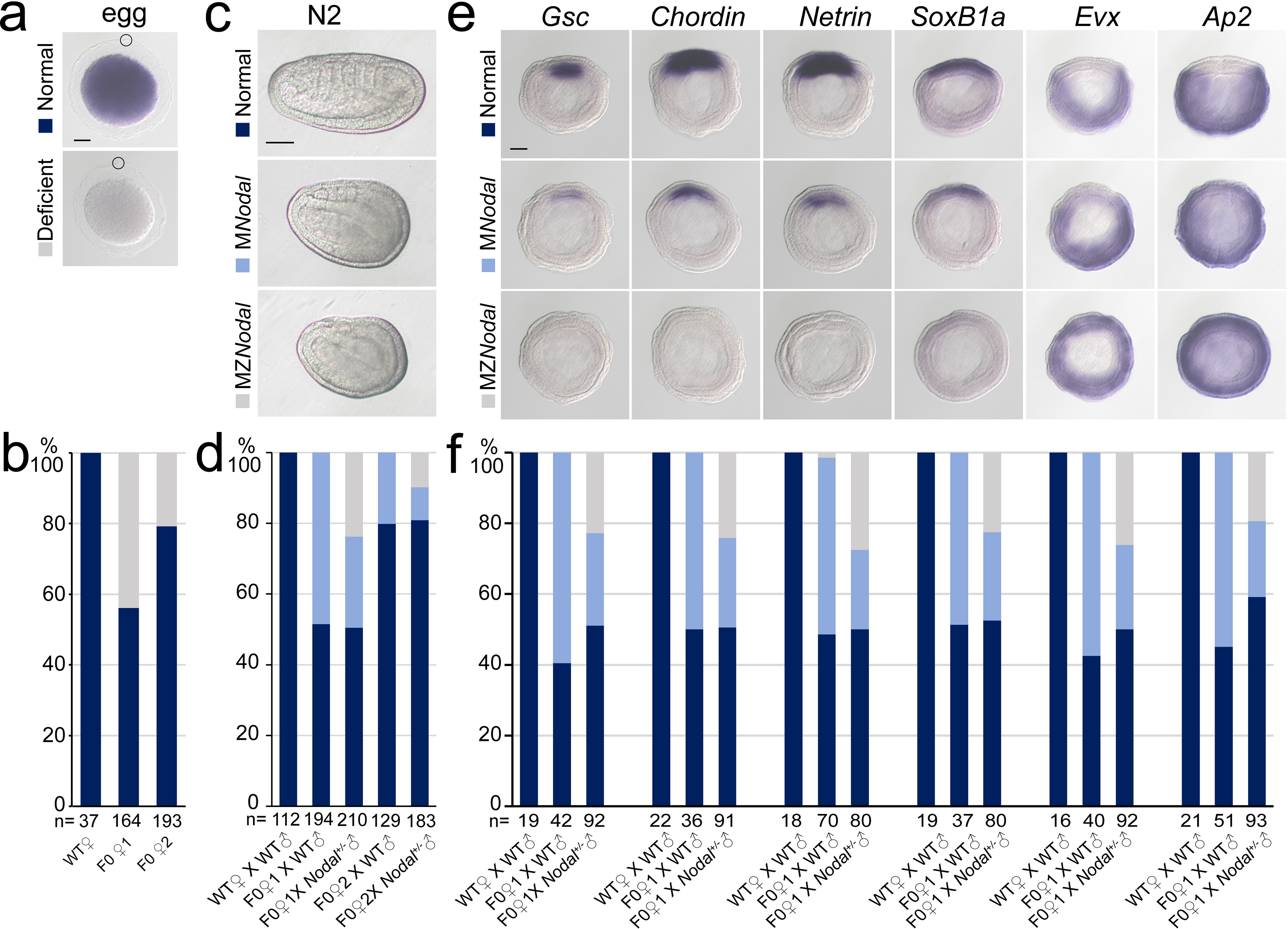
*Nodal* gene is required for amphioxus axis development. **(a)** *Nodal* expression in amphioxus eggs from different females (WT, F0♀1 and F0♀2). They were observed with animal pole to the top (circles indicate polar body). Two types of *Nodal* expression (normal and deficient) were observed. **(b)** Histogram showing the percentage of eggs of normal or deficient maternal *Nodal* accumulation from different females. **(c)** Phenotypic analyses of maternal (M*Nodal*) and maternal-zygotic (MZ*Nodal*) *Nodal* mutants at N2 stage. They were observed from the left side with anterior to the left at N3 stage. Phenotype of the two mutants at other stages are provided in Supplemental Figure 7. **(d)** The percentage of embryos of different phenotypes as shown in (c). Embryos from five different crosses were examined. **(e)** The expression of marker genes in M*Nodal* and MZ*Nodal* mutants at G5 stage. All embryos were viewed from the blastopore with dorsal to the top. **(f)** The percentage of embryos showing different expression patterns (normal, M*Nodal* and MZ*Nodal*) as indicated in (e). Embryos from three different crosses were examined. Scale bars, 50 μm (a, c and e). The total number of analyzed eggs or embryos are listed under each column (b, d and f).

The phenotypes of MZ*Nodal* and *Nodal^-/-^* are strikingly similar to those of *Gdf1/3-like^-/-^* and *Gdf1/3-like^+/-^* respectively, implying that Nodal is indispensable for Gdf1/3-like activity during Nodal signal transduction. To further test this, we injected either *Nodal* or *Gdf1/3-like* mRNA into maternal M*Nodal* embryos and analyzed the *Gsc* expression at the G5 stage. We found that *Nodal* mRNA injection could rescue *Gsc* expression in the M*Nodal* embryos, while *Gdf1/3-like* mRNA injection could not, although it expanded the *Gsc* expression domain in normal embryos (Supplementary Fig. 9). This result demonstrated that Gdf1/3-like alone (without Nodal) is unable to activate Nodal signaling in amphioxus.

### *Gdf1/3-like* hijacked the regulatory region of *Lefty*

As *Gdf1/3-like* is linked to *Lefty* in a head-to-head way in all sequenced amphioxus genomes and the two genes exhibit a similar expression pattern during embryogenesis, we hypothesized that the intergenic region between the two genes might include most if not all cis-regulatory elements (such as enhancers) required for their expression. To test this, we cloned the intergenic region (about 4 kb) into a pminiTol2 plasmid carrying a *mCherry* reporter and generated a stable amphioxus transgenic line carrying it (Fig. 4a). Whole mount *in situ* hybridization analysis revealed that the transcription of *mCherry* was similar to that of endogenous *Gdf1/3-like* and *Lefty* with symphonic initiation and D-V and L-R asymmetric expression pattern (Fig. 4a). We further made a dual-reporter pminiTol2 construct, in which the coding sequences of *eGFP* and *mCherry* were inserted into the two ends of the 4 kb region respectively, and injected it into amphioxus embryos with mRNAs of Tol2 transposase. *In situ* analysis showed that at the G0 stage, the 4 kb sequence could simultaneously drive *eGFP* and *mCherry* transcription in a similar pattern in the dorsal organizer region (Fig. 4b). These results indicated that the intergenic sequence indeed contains sequence elements required for regulating the expression of the two linked genes.

**Figure 4.**
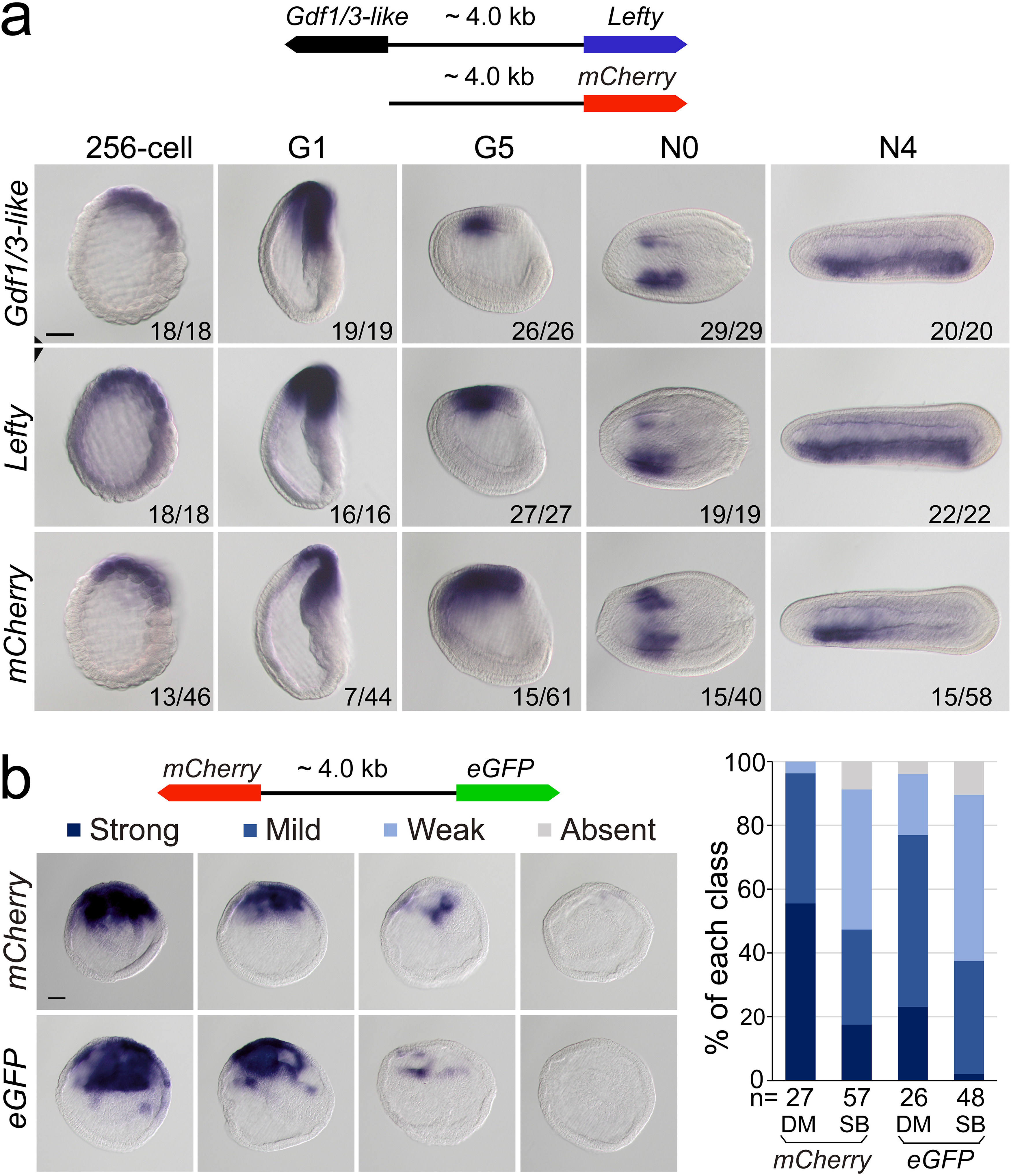
Regulatory activity of the intergenic region between *Gdf1/3-like* and *Lefty* genes. (a) The top schematic diagram shows the 4 kb region between *Gdf1/3-like* and *Lefty* and the construct used to generate stable amphioxus transgenic lines. Embryos used for *in situ* analysis were from a cross between a female founder and a WT male. The bottom panels show the expression patterns of *Gdf1/3-like*, *Lefty* and *mCherry* reporter in transgenic embryos. Embryos at 256-cell to G5 stages were viewed from the left side, and those at N0 and N4 stages were viewed from the dorsal side. All embryos are anterior to left. Numbers at the bottom right of each panel indicate the number of times the expression pattern was observed, out of the total number of examined embryos. (b) The left top schematic diagram shows the dual-reporter construct used in this analysis, and the panels below it shows the expression of *mCherry* and *eGFP* in embryos injected with the construct and then treated with DMSO (DM) or SB505124 (SB). All embryos are at G1 stage viewed form the blastopore with dorsal to the top. Four categories of expression (strong, mild, weak and absent) for both *mCherry* and *eGFP* were observed in the dorsal blastopore of embryos, and their percentages are shown in the right histogram. The total number of analyzed embryos are listed under each column. Scale bars, 50 μm.

After demonstrating the bidirectional activity of the 4 kb intergenic sequence in driving *Lefty* and *Gdf1/3-like* genes, we then asked which one of the two genes originally used this region to regulate their expression. In sea urchin and vertebrate embryos, the expression pattern of *Lefty* and *Gdf1/3* genes are different, with the former being zygotically activated at blastula stage and unilaterally expressed from gastrula/neurula stage^22-28^, and the latter being maternally preloaded and bilaterally expressed from gastrula/neurula stage (Supplementary Table 2). Moreover, *Lefty* expression depends on Nodal signaling in these two groups^29,30^, while *Gdf1/3* expression does not, at least in sea urchin^19^. As demonstrated above, *Lefty* expression pattern in amphioxus is similar to its cognates in sea urchin and vertebrates, but *Gdf1/3-like* expression follows essentially that of *Lefty*. Additionally, we and others also showed that *Lefty* expression in amphioxus embryos depends on Nodal signaling^9-12^. These results therefore implied that the 4 kb region was initially responsible for *Lefty* expression, and then hijacked by *Gdf1/3-like* after it was translocated to the current locus in amphioxus ancestor. To evaluate this scenario further, we examined *Gdf1/3-like* expression in the M*Nodal* embryos to see if it is dependent on Nodal signaling as that of *Lefty*. The result showed that, in M*Nodal* embryos, the expression of *Gdf1/3-like* and *Lefty* were both significantly reduced or abolished at the gastrula stages (G1-G5), although their initial expressions at the blastula stage (256-cell) were not affected (Supplementary Fig. 10). To examine this more directly, we injected amphioxus embryos with the above dual-reporter construct and then treated them with Nodal signaling inhibitor SB505124 to see if the expression patterns of both reporter genes are affected. *In situ* result showed that compared to untreated embryos, SB505124-treated embryos exhibited decreased expression for both *eGFP* and *mCherry* genes (Fig. 4b). This indicates that the 4 kb region includes the sequence elements required for *Lefty* and *Gdf1/3-like* expression regulation by Nodal signaling. Together, these results imply that the 4 kb region contains regulatory elements which were initially used to regulate *Lefty* expression in different deuterostomes and later hijacked by *Gdf1/3-like* after its translocation next to *Lefty* gene in amphioxus.

## Discussion

During evolution, morphological innovations are usually accompanied by changes of GRNs essential for developmental processes^2^. However, the GRNs could also evolve or diverge without altering homologous characters, a phenomenon called developmental system drift^31^. In either way, functional genetic evidence with a robust phylogenetic framework to demonstrate how GRN changes could have happened is sparse so far, although the underlying mechanism for the evolvability has been usually attributed to changes of cis-regulatory elements^32^. The most striking discovery in our study is that enhancer hijacking events could trigger the evolution of a GRN. In addition, through detailed analyses of available functional genetic data below, we could demonstrate how a GRN could have evolved in a stepwise way, in the absence of “molecular developmental fossils” (Fig. 5).

**Figure 5.**
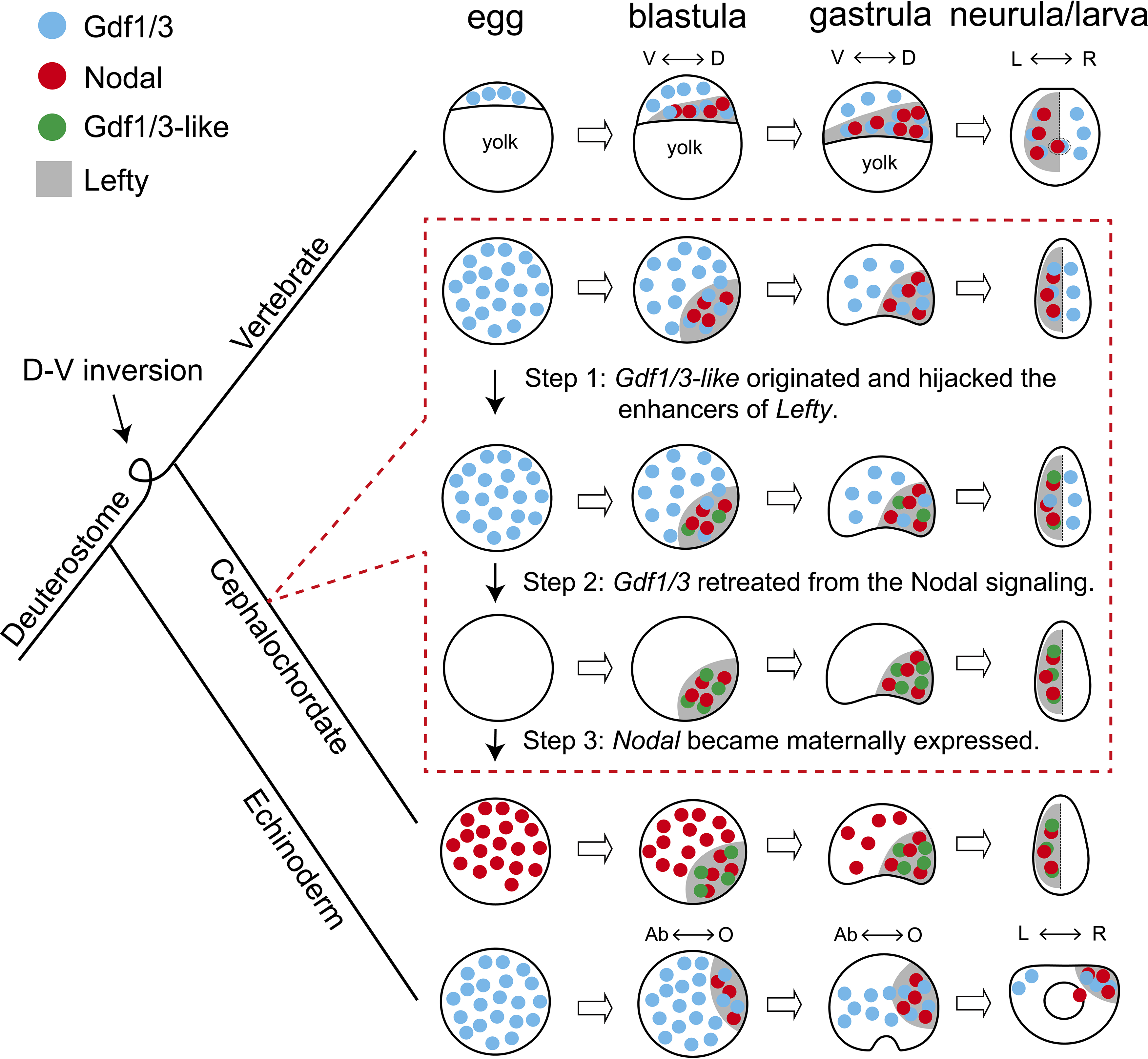
Scenario for evolution of the Nodal signaling in amphioxus. The situation in sea urchin (echinoderm) and zebrafish (vertebrate) represents an ancestral scenario which has been rewired and modified in amphioxus through at least three sequential steps as shown in dashed box. D-V, dorsal-ventral axis. L-R, left-right axis. Ab-O, oral-aboral axis. The dorsal-ventral orientation of chordates was inversed relative to the ambulacraria during evolution (Lowe et al., 2015), thus the oral and left side of sea urchin corresponds to the dorsal and right side of zebrafish and amphioxus respectively.

Nodal signaling plays an essential role for patterning the D-V and L-R axis in deuterostomes^3^. The expression patterns of the genes coding the ligands of the signaling pathway are conserved in extant echinoderms and vertebrates (Supplementary Table 1 and 2). Thus, the GRN of the Nodal signaling pathway revealed in these species most likely represents an ancestral scenario: Gdf1/3 is preloaded in an egg; by blastula stage, Nodal and Lefty appear zygotically; Nodal functions interdependently with the preexisting Gdf1/3 to activate the signaling pathway, while Lefty restrains the activity of the signaling pathway by inhibiting the signaling (Fig. 5)^4,6^. Our data showed that this highly conserved GRN has been rewired and modified in cephalochordate amphioxus: Nodal is expressed maternally in an egg; ancestral Gdf1/3 lost its expression and function at embryonic stage, while *Gdf1/3-like* (a duplicate of *Gdf1/3*) recruited the regulatory elements of *Lefty* and is expressed in a similar pattern as *Lefty* from the blastula stage; Gdf1/3-like functions interdependently with the preexisting Nodal to activate the signaling pathway; and Lefty still acts as an feedback inhibitor of the signaling pathway (Fig. 5). Our results and those from previous studies in other species enable us to infer that at least three sequential steps are probably required to evolve the GRN as found in extant amphioxus from an ancestral situation as reported in echinoderms and vertebrates (as discussed below): 1. the *Gdf1/3-like* originated and translocated adjacently to *Lefty* and hijacked its enhancers; 2. the *Gdf1/3* retreated from the Nodal signal; 3. the *Nodal* became maternally expressed (Fig. 5).

Overexpression of *Gdf1/3* in zebrafish and *Xenopus* embryos have no effects on axis development^30,33^. It is therefore expected that the emergence of *Gdf1/3-like* would not affect the development and viability of the ancestor of extant amphioxus (i.e. step 1, Fig. 5). By contrast, disruption of maternal *Gdf1/3* function is lethal in sea urchin and zebrafish embryos^4,19,30,34^. This indicates that the retreatment of the *Gdf1/3* orthologue from the Nodal signaling (step 2) could not happen before the emergence of *Gdf1/3-like* and its recruitment of the *Lefty* enhancers (step 1). Similarly, overexpression of Nodal in the presence of maternal Gdf1/3 led to defects in axis formation of zebrafish embryos^30^, suggesting that Nodal could not become a maternal factor before the retreatment of *Gdf1/3* (i.e. step 2 happened before step 3). These suggested that the 1-2-3 ordered events could have led to the situation in extant amphioxus (Fig. 5). Injection of *Gdf1/3* mRNA to the yolk syncytial layer, where *Nodal* is expressed endogenously, was sufficient to rescue M*Gdf1/3* defects in zebrafish, thus the ubiquitous distribution of maternal *Gdf1/3*, in the region where *Nodal* is not expressed, is dispensable^4^. Once the *Gdf1/3-like* started its current expression pattern by hijacking *Lefty* enhancers, the function of *Gdf1/3-like* and *Gdf1/3* became redundant, allowing the retreatment of *Gdf1/3* from the signaling pathway (i.e. step 2, Fig. 5). As discussed in the study of zebrafish^4^, preloading of a maternal factor (Gdf1/3 in zebrafish or Nodal in amphioxus) could be instrumental for ensuring Nodal signaling initiation in a rapid and temporally reliable manner. Likewise, the establishment of Nodal as a maternal factor may be a compensation mechanism for the retreatment of ubiquitously maternal Gdf1/3 (i.e. step 3, Fig. 5). Zygotic *Nodal* mutants failed to form L-R axis but formed D-V axis normally in amphioxus (Supplementary Fig. 6), suggesting that maternal *Nodal* is redundant to compensate the D-V axis formation. However, maternal *Nodal* mutants formed only partial anterior and dorsal structures (Fig. 3, Supplementary Fig. 7 and 8), suggesting that the expression of zygotic *Nodal* might be delayed in living amphioxus, compared to that of the amphioxus ancestor before step 3. Our proposed scenario highlights that enhancer hijacking by *Gdf1/3-like* was a triggering and probably “neutral” step for the rewiring of the body axes GRN in amphioxus ancestors.

Although developmental system drift is probably universal in long evolutionary periods^35^, it has been rarely possible to analyze whether there is selective advantage of such drifts. Co-expressed gene pairs sharing intergenic enhancers have been reported in various cases^36-39^, and often fall into the same functional categories^38^. In zebrafish, the paired *her1* and *her7* can provide robustness for segmentation patterning^39^. The two proteins need to form dimers to inhibit their own transcription, thus forming a negative-feedback loop to maintain a stable level of both proteins in cells^39^. In our case, Gdf1/3-like functions as an activator of the signaling that maintains the transcriptions of both *Gdf1/3-like* and *Lefty* in a positive-feedback loop, while Lefty acts as a repressor of the signaling forming a negative-feedback loop^13,14^. The co-expression of *Gdf1/3-like* and *Lefty* gene pair (Fig. 1b, Supplementary Fig. 1) achieved by sharing regulatory region (Fig. 4) likely safeguards a dosage balance between the positive and negative-feedback loop. This is likely an advantage for robust pattern formation as suggested in other gene pairing systems^38^.

Amphioxus has been shown to possess small-scale gene duplications^40^, and *Gdf1/3* is one such example. Interestingly, *Gdf1/3* has also been duplicated independently in anurans and mammals^17^. However, these genes appear to have evolved differently: in anurans and mammals both *Gdf1/3* paralogs retain the ancestral function in axis formation^41-43^, while in amphioxus only *Gdf1/3-like* undertakes this function but *Gdf1/3* seems to have lost its original function. As discussed above, hijacking of *Lefty* enhancers by *Gdf1/3-like* had enabled *Gdf1/3-like* to be functionally redundant to *Gdf1/3*, allowing *Gdf1/3* to retrieve from its ancestral function in regulating body axis formation through changing its expression pattern. In addition, the redundancy of *Gdf1/3-like* appears to have also relaxed the selection pressure on *Gdf1/3* coding region, since *Gdf1/3-like* mRNA is more efficient in inducing anterior and dorsal expansion than that of *Gdf1/3* (Supplementary Fig. 3).

We have demonstrated that functional genetic data could be used to dissect the route of GRNs evolution, once there is available data from different species for comparisons. Our study represents an example that illustrated how the GRNs could have shifted in a stepwise way without altering conserved phenotypes, such as body axes that has persisted at least since the Early Cambrian period^44,45^. Our case also showed that enhancer hijacking could be a mechanism underlying the evolution of developmental GRNs, in addition to the changes of cis-regulatory elements.

## Methods

### Animals and embryos

Wild type (WT) amphioxus (*Branchiostoma floridae*) was obtained from Jr-Kai Yu’s lab and bred in the aquaculture system as reported previously^46^. Mature individuals were induced to lay eggs and release sperm by thermal shock^47^. The developmental stages of amphioxus embryos were defined as recently described^48^.

### Gene structures and synteny

Positions of *Lefty*, *Gdf1/3-like*, *Bmp2/4* and *Gdf1/3* genes in the genomes of *B. floridae*, *B. belcheri*, *B. japonicum* and *B. lanceolatum* were determined according to two recent studies^40,49^, while positions of these genes in sea urchin (*S. purpuratus*) were retrieved directly from Echinobase, in other species including zebrafish *(D. rerio*), elephant Shark (*C. milii*), vase tunicate (*C. intestinalis*), acorn worm (*S. kowalevskii*), polychaete worm (*C. teleta*) and fruit fly (*D. melanogaster*) were retrieved directly from Ensembl database.

### Quantitative real-time PCR

Embryos or eggs (about 200 per sample) were harvested at desired stages and used to extract total RNA by TRIzol reagent (Ambion). cDNA was synthesized using HiScript III RT SuperMix (+gDNA wiper) kit (Vazyme). Quantitative real-time PCR analysis was performed with ChamQ Universal SYBR qPCR Master Mix kit (Vazyme) on a CFX96 Touch Real-Time PCR Detection System (Bio-Rad) under the conditions of 95 °C for 2 min, 40 cycles at 95 °C for 5 s, 60 °C for 30 s. Expression levels of *Nodal*, *Gdf1/3*, *Gdf1/3-like* and *Lefty* were normalized to that of *Gapdh* (glyceraldehyde-3-phosphate dehydrogenase). Graphs were finished with GraphPad Prism 9 software. Primers for qRT-PCR analysis and their and sequences were as follows: BfGdf1/3-like-RT-F (5′-CAAGGGCAAATATCACGACA-3′), Bf Gdf1/3-like-RT-R (5′-TTCACGTCGTCTCTGTCGAA-3′), BfNodal-RT-F (5′-GGACAGACCTCAACGTCACCC-3′), BfNodal-RT-R (5′-CTGAAGACACGCACGGAAAGT-3′), BfLefty-RT-F (5′-CACTGACGCCAGTGGTGCA-3′)、BfLefty-RT-R (5′-CGTTGTTGAAAGACTTTCGAGT-3′), BfGdf1/3-RT-F (5′-TTCTCGGCTTTCGTGAACGG-3′), BfGdf1/3-RT-R (5′-ACAGTCCAACCATTTTCGGCA-3′), Gapdh-RT-F (5′-GGTGGAAAGGTCCTGCTCTC-3′), and Gapdh--RT-R (5′-CTGGATGAAAGGGTCGTTAATGG-3′).

### Overexpression experiments

Coding sequences of *Nodal* and *Gdf1/3-like* were amplified from cDNA templates of amphioxus embryos and ligated into the pXT7 vector using T4 DNA ligase (Promega). Due to low expression level, coding exons of *Gdf1/3* were individually amplified from genomic DNA templates and then assembled into the pXT7 vector using a Gibson cloning kit (New England Biolabs). *Nodal*, *Gdf1/3* and *Gdf1/3-like* mRNA were synthesized using T7 mMESSAGE mMACHINE kit (Ambion). Unfertilized eggs were injected with mRNAs and fertilized as previously described^50^. The injected embryos were treated, with 0.2% DMSO (as control) or 50 μM SB505124, from 4-cell stage to G1 stage and were fixed at G4 or N3 stage for whole mount *in situ* hybridization.

### Generation of mutant animals and embryos

*Gdf1/3* mutant were generated using CRISPR/Cas9 system targeting a site (*Gdf1/3-sg*RNA: 5’-GGCCCGCTGTAGCGATGA-3’) in the first exon. The process of generating founders of *Gdf1/3* gene knock-out was implemented as our previous study^51^. F1 *Gdf1/3^+/-^* carrying -25 bp mutation were intercrossed to generate *Gdf1/3 ^-/-^* (Supplementary Fig. 2). A pair of primer (*Gdf1/3*-Cas9-F1: 5’-TACCACACATCACCCGGACT-3’/*Gdf1/3*-PCR-R1: 5’-CACATCCTCGTCTTCCGGTC-3’) and *Sfc*I enzyme were used for genotyping and mutation type analysis.

*Gdf1/3-like* mutants were generated with a TALEN pair (*Gdf1/3-like*-Fw: 5’-TTCGACAGAGACGAC-3’/*Gdf1/3-like*-Rv: 5’-TGCACGGCGCTCACGA-3’) targeting the coding region of the second exon as previously reported^13,52^. F1 heterozygotes carrying a one base pair deletion (+1 bp) were used and intercrossed to generate homozygote mutants (*Gdf1/3-like^-/-^*) (Supplementary Fig. 4). Embryos or tiny tips of adult tail were lysed with Animal Tissue Direct PCR Kit (Foregene) to release the genomic DNA. Genomic region spanning the target site were amplified with primer *Gdf1/3-like*-TALEN-F: 5’-CGTGACGTACTCCGTGTCTG-3’/*Gdf1/3-like*-TALEN-R1: 5’-GCTGAAGTGTGGGCAAGAGT-3’or *Gdf1/3-like*-TALEN-R0: 5’-CCGTTTGCAGATGTTGCCG-3’. The amplicons were digested with *Stu*I to test the mutations and sequenced to recognize the mutation types.

*Nodal* heterozygotes (*Nodal^+/-^*) were generated using a TALEN pair (Nodal-Fw2/Rv2) in previous study^53^. Male and female *Nodal^+/-^* carrying identical mutation (-7 bp) were intercrossed to generate *Nodal^-/-^* (Supplementary Fig. 6). Genotyping for *Nodal* mutants were implemented as previous study^53^. A TALEN pair (Nodal-Fw2/Rv2) assembled previously^53^ and a sgRNA (*Nodal*-sgRNA:5’-GGCGGAGAGGGTCTGACGCT-3’) reported previously^51^ were used to generate hundreds of chimeric female founders. By adulthood stage, among nearly one hundred of animals, an individual generated with Nodal-Fw2/Rv2 (named founder 1, F0♀1) and another individual generated with *Nodal*-sgRNA (named founder 2, F0♀2) laid eggs lacking maternal *Nodal* accumulation (Supplementary Fig. 7a) with 50% and 20% ratio respectively. Each female founder was crossed with a male WT respectively to generated maternal *Nodal* mutants (M*Nodal*) and crossed with a male *Nodal^+/-^* respectively to generated maternal-zygotic *Nodal* mutants (MZ*Nodal*) (Supplementary Fig. 7b).

### Generation of transgenic animals and embryos

The intergenic region between *Gdf1/3-like* and *Lefty* was cloned into the pmini-Chordin-mCherry^54^ by replacing its *Chordin* promoter with the intergenic region. The generated pmini-Lefty-mCherry construct was used to generate *Lefty::mCherry* transgenic amphioxus with method reported previously^54^. The intergenic region, the coding sequence of *mCherry* and *eGFP* sequence were linked into the pminiTol2 plasmid with a Gibson cloning kit (New England Biolabs) to generate pmini-eGFP-Lefty-Gdf1/3-like-mCherry construct. The later was injected into amphioxus embryos. The injected embryos were treated, with 0.2% DMSO (as control) or 50 μM SB505124, from 4-cell stage to G1 stage.

### Whole mount *in situ* hybridization and imaging

The RNA probes used in present study were prepared previously^13,14,54^, except the probe of *eGFP* and *Gdf1/3* gene, which was prepared using the same method as described in a previous study^13^. Embryos at desired developmental stages were fixed in 4% PFA-MOPS (wt/vol) and then stored in 80% ethanol (vol/vol) in H2O at -20°C until needed. Whole-mount *in situ* hybridization was performed as previously described ^55^. After staining, the embryos were mounted in 80% glycerol (vol/vol) for photographing under an inverted microscope (Olympus IX71). Living amphioxus embryos or larvae were observed using the same microscope.

## Data availability

Pre-processed data and other findings of this study are available from the corresponding authors upon reasonable request.

## Acknowledgements

We thank all team members of the G.L. and Q.Q. laboratories, and Prof. Zhenghong Zuo for helpful discussions. This work is supported by grants from the National Natural Science Foundation of China (nos. 32070458, 31872186, 32070815 and 32061160471), and from the Youth Innovation Fund Project of Xiamen (3502Z20206032).

## Author contributions

G.L., Q.Q., C.S. and Y.W. designed the experiments. C.S., S.C., X.L., C.X., X.L. and G.L. generated the mutant and transgenic lines. C.S., S.C., H.L., S.L., Y.W., R.P. and X.W. performed the whole mount *in situ* hybridization, phenotype analyses, genotyping and over expression experiments. S.L. performed the quantitative real-time PCR. G.L., J.L. and C.S. carried out the genome survey. C.S., G.L. and Q.Q. interpreted the data, prepared the figures and wrote the manuscript. All authors commented on the manuscript and agreed to its final version.

## Competing interests

The authors declare no competing interests.

---

**Supplementary Figure 1.**
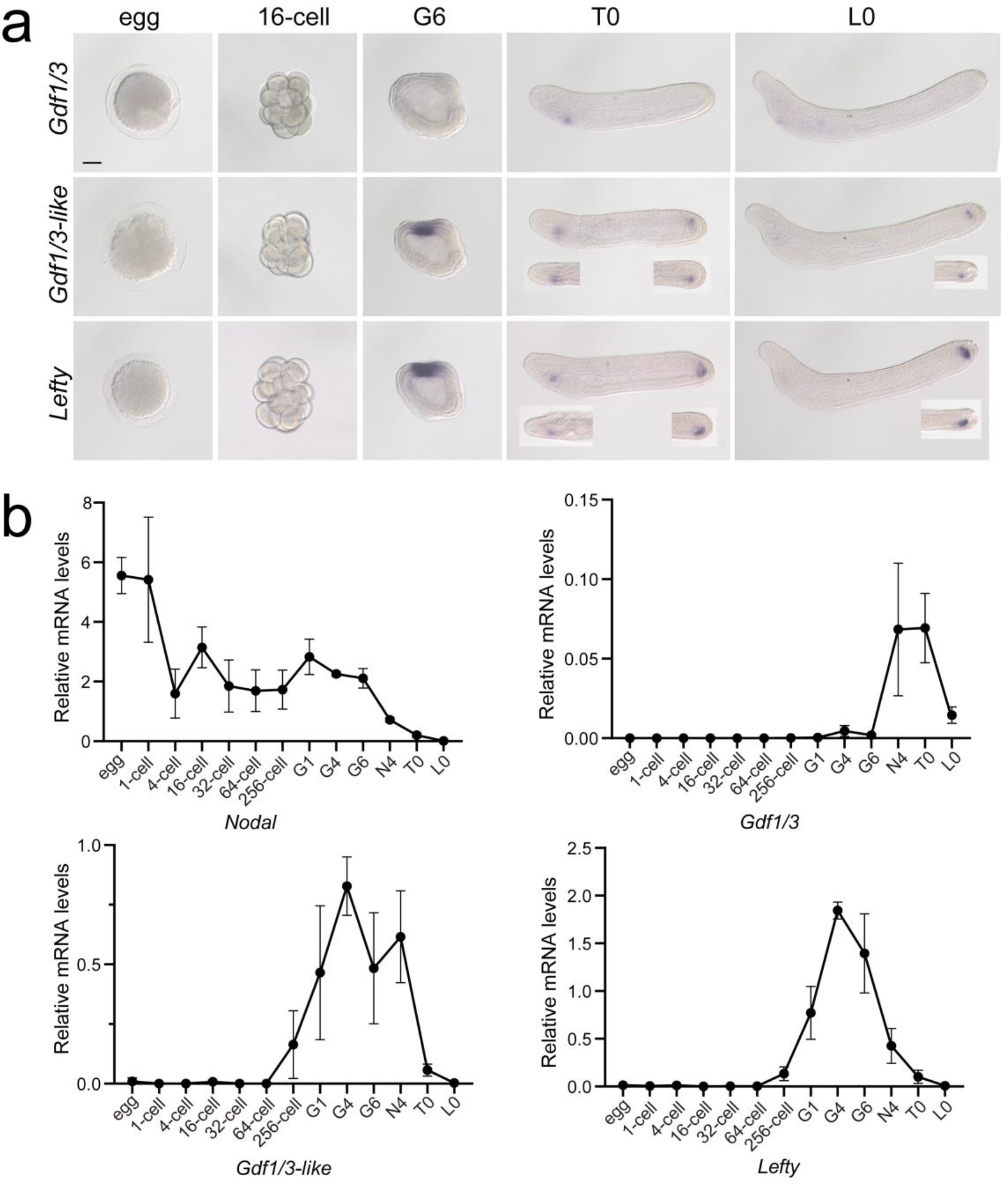
Analysis of *Gdf1/3*, *Gdf1/3-like* and *Lefty* expression patterns in amphioxus embryos at different stages. **(a)** Embryos, analyzed with *in situ* hybridization, at 16-cell to L0 stage were viewed from the left side with anterior to the left. Inserts in the panels of some T0 or L0 embryos were viewed from the dorsal side. Scale bar, 50 μm. **(b)** The expression level of examined genes, analyzed with quantitative real-time PCR, was normalized to that of *Gapdh*. Error bars indicate standard deviation from 3 biological replicates.

**Supplementary Figure 2.**
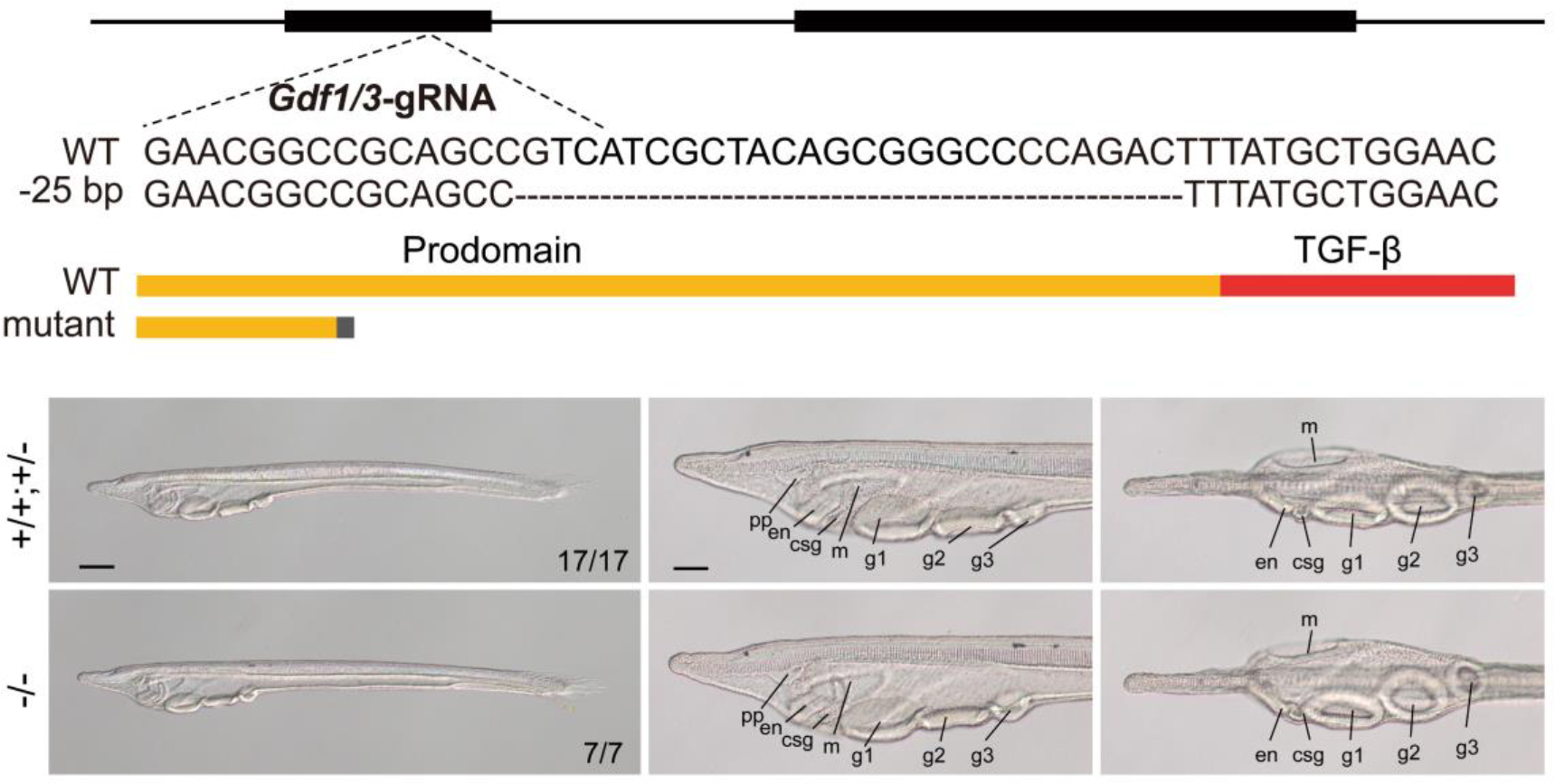
Phenotypic analysis of amphioxus *Gdf1/3* mutants. The top schematic diagram shows the exon/intron structure of *Gdf1/3* coding sequence, the sgRNA used to target it, the introduced mutation (a 25-bp deletion), and the structure of the truncated proteins encoded by the mutated *Gdf1/3* gene. Black boxes indicate *Gdf1/3* exons. Panels in the bottom left show whole mount of *Gdf1/3* larvae (L3 stage) of different genotypes (*+/+*, *+/-* and -/-) observed from the left side with anterior to the left. Numbers in the panels indicate the number of times the genotype shown was identified, out of the total number of larvae identified with the phenotype. Panels in the bottom middle and right are anterior parts of *Gdf1/3* larvae respectively observed from the left side and ventral side, with anterior to the left. pp, preoral pit; en, endostyle; csg, club shaped gland; m, mouth; g1, first gill slit; g2, second gill slit; g3, third gill slit. Scale bars in panels on the left are 100 μm, and those in other panels are 50 μm.

**Supplementary Figure 3.**
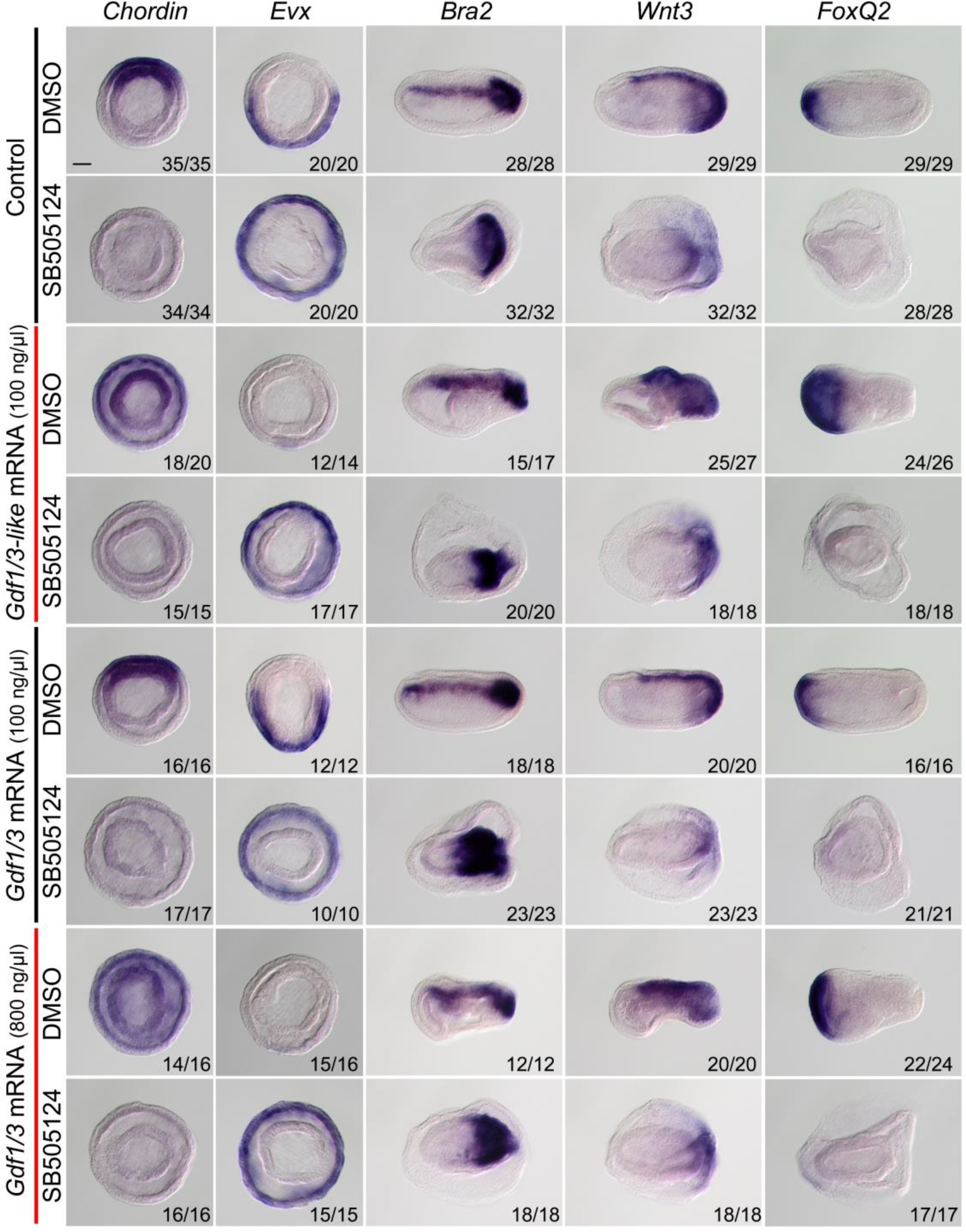
The expression of marker genes in embryos injected with *Gdf1/3* or *Gdf1/3-like* mRNA and treated with DMSO or SB505124. Embryos stained with *Chordin* and *Evx* probes were at G4 stage and observed from the blastopore with dorsal to the top. Embryos stained with *Brachyury*, *Wnt3* and *FoxQ2* probes were at N3 stage and observed from the left side with anterior to the left. Numbers at the bottom right of each panel indicate the number of times the expression pattern shown was observed, out of the total number of examined embryos. Scale bar, 50 μm.

**Supplementary Figure 4.**
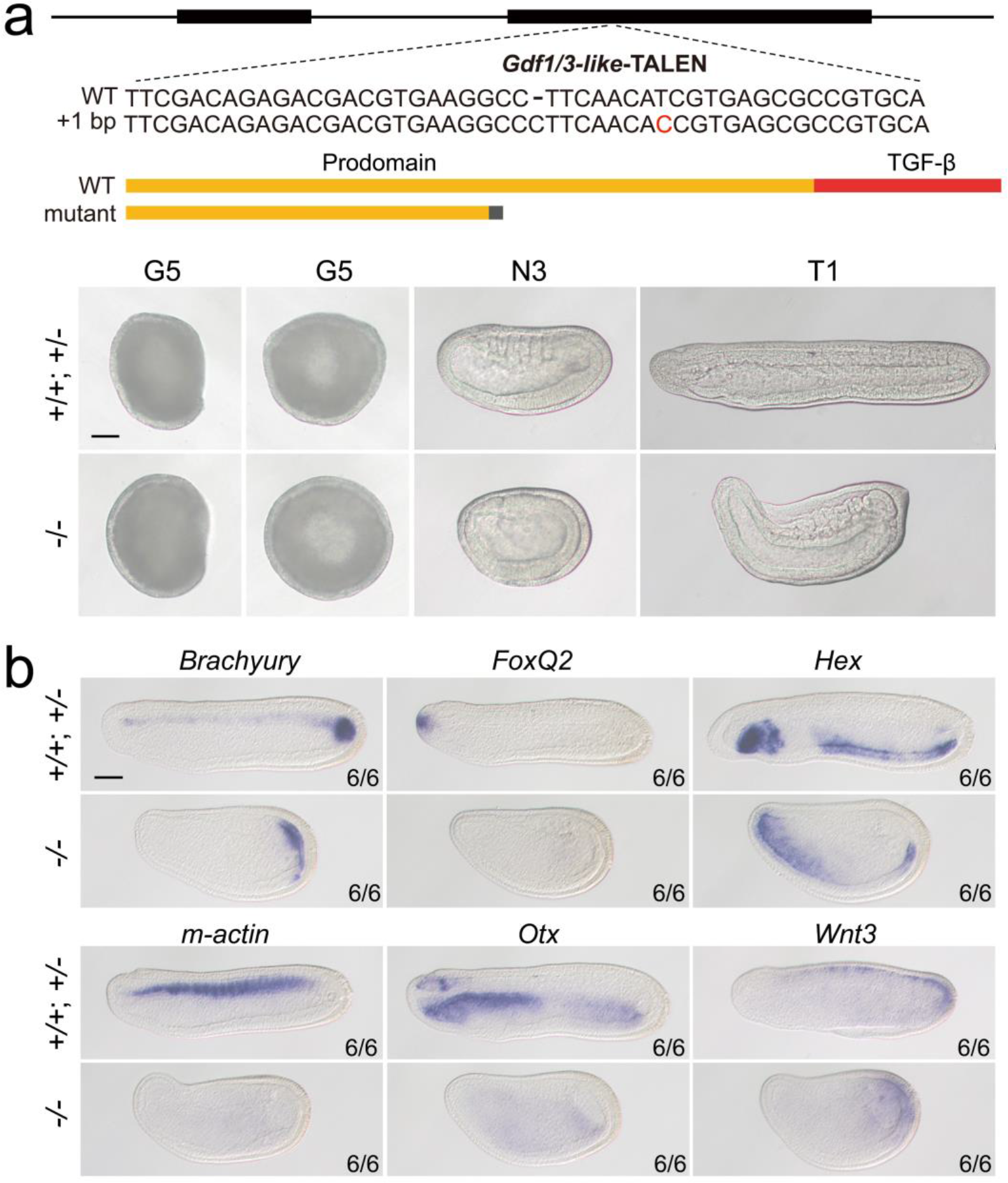
Phenotypic analysis of *Gdf1/3-like* mutants and the expression of marker genes in *Gdf1/3-like* mutants. **(a)** The top schematic diagram shows the exon/intron structure of *Gdf1/3-like* coding region, the TALEN used to target it, the introduced mutation (a 1-bp insertion), and the structure of the truncated proteins encoded by the mutated *Gdf1/3-like* gene. Black boxes indicate exons. Panels in the bottom show *Gdf1/3-like* embryos of different genotypes (*+/+*, *+/-* and -/-). Embryos at G5 stage were observed from either the left side with anterior to the left (left ones), or from the blastopore with dorsal to the top (right ones). Embryos at other stages were observed from the left side with anterior to the left. **(b)** All embryos were observed from the left side with anterior to the left at T0 stage. Numbers at the bottom right indicate the number of times the genotype shown was identified, out of the total number of larvae identified with the phenotype or expression pattern. Scale bars, 50 μm.

**Supplementary Figure 5.**
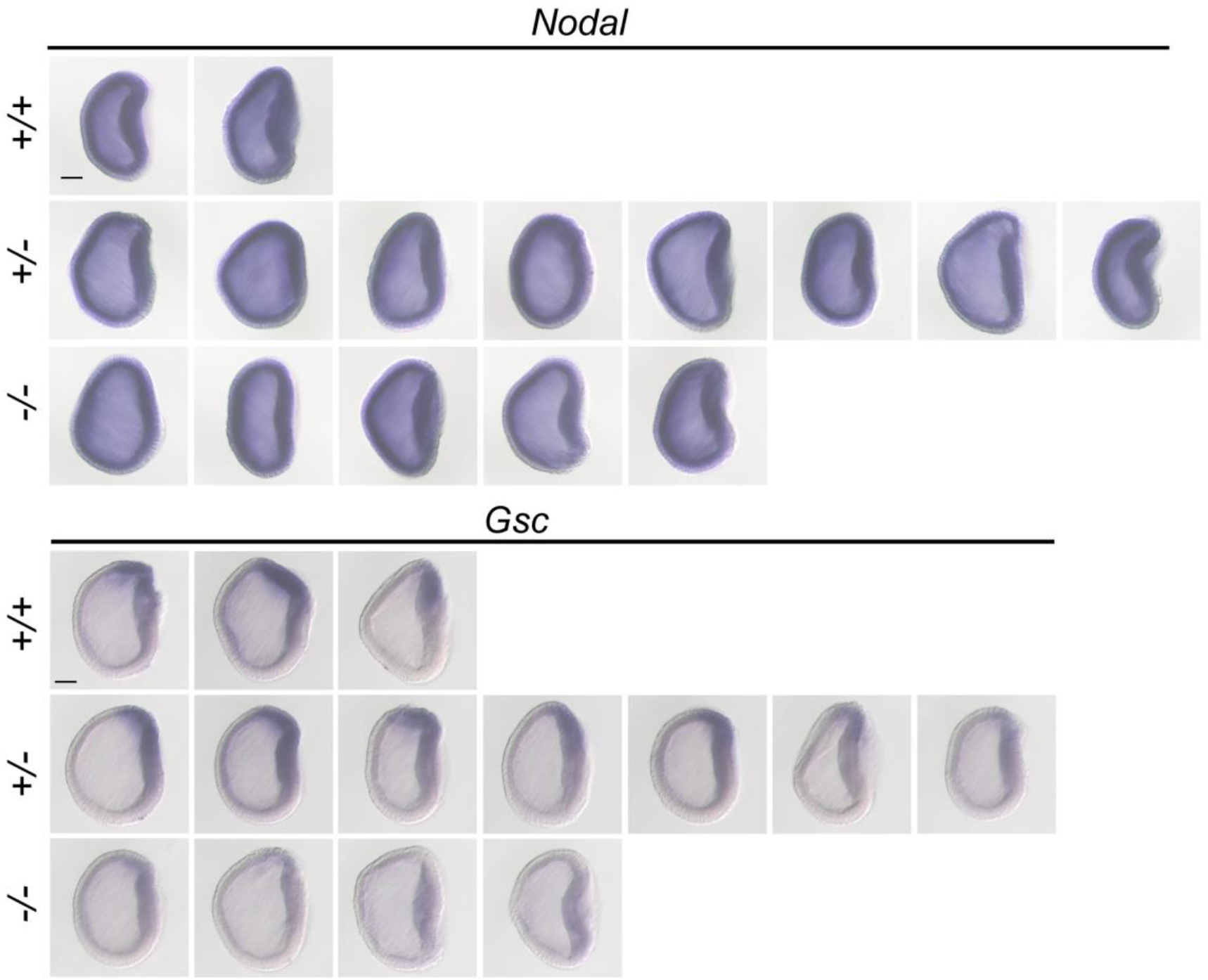
The expression pattern of *Nodal* and *Gsc* in *Gdf1/3-like* mutants at G1 stage. All embryos identified with each genotype (*+/+*, *+/-* and -/-) in the analysis were shown and they were all viewed from the left side with anterior to the left. Scale bars, 50 μm.

**Supplementary Figure 6.**
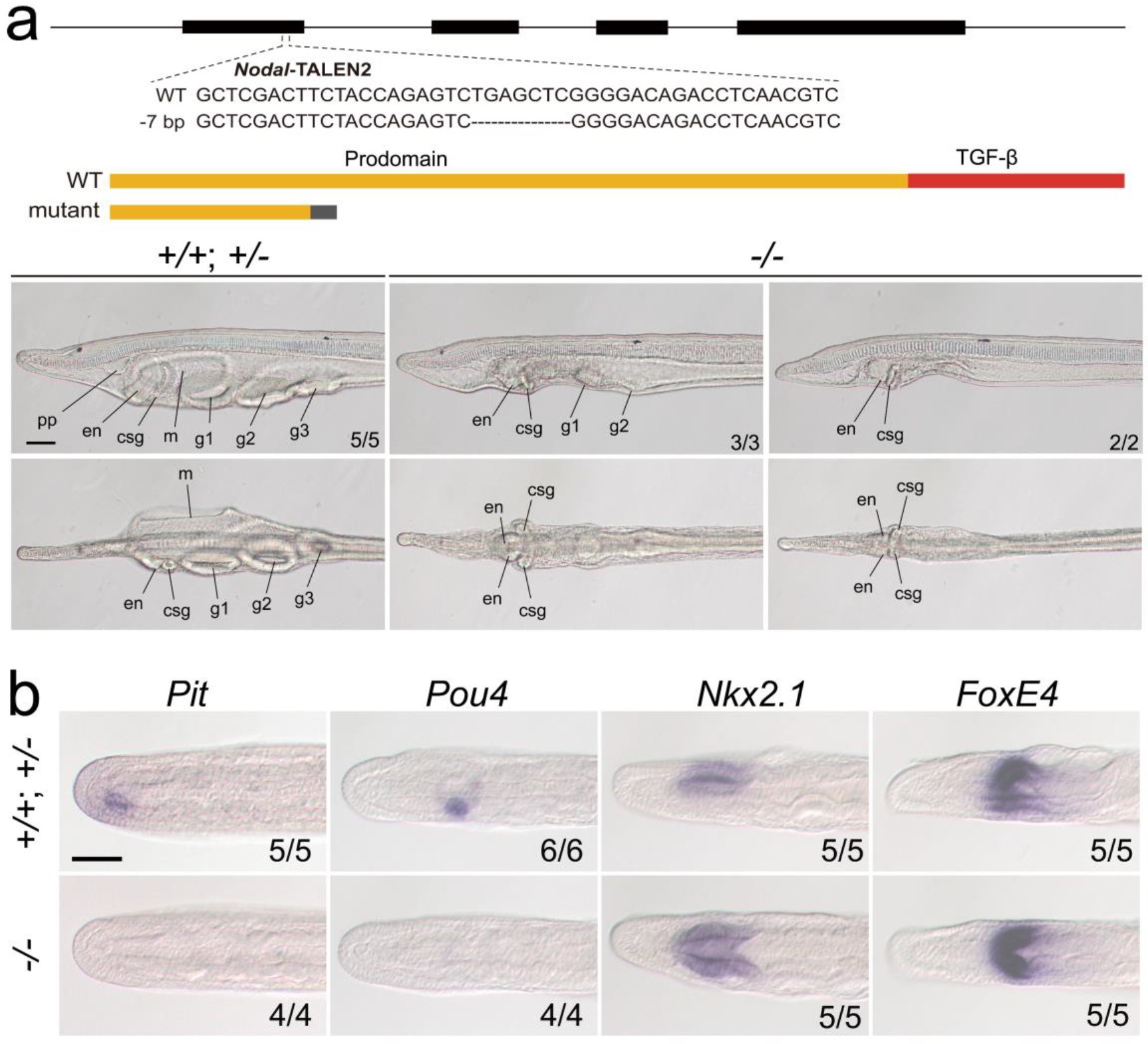
Phenotypic analysis of zygotic *Nodal* mutants and the expression of marker genes in zygotic *Nodal* mutants. **(a)** The top schematic diagram shows the exon/intron structure of *Nodal* coding sequence, the TALEN used to target it, the introduced mutation (a 7-bp deletion), and the structure of the truncated proteins encoded by the mutated *Nodal* gene. Black boxes indicate exons. Panels in the bottom show the anterior part of *Nodal* larvae (L3 stage) of different genotypes (*+/+*, *+/-* and -/-), observed from the left side (upper panels) and ventral side (lower panels) with anterior to the left. Numbers at the bottom right indicate the number of times the genotype shown was identified, out of the total number of larvae identified with the phenotype. pp, preoral pit; en, endostyle; csg, club shaped gland; m, mouth; g1, first gill slit; g2, second gill slit; g3, third gill slit. **(b)** All panels are anterior parts of embryos observed from the dorsal side at T0-T1 stage, with anterior to the left. Numbers at the bottom right indicate the number of times the genotypes shown were identified, out of the total number of examined larvae with the expression pattern. Scale bars, 50 μm.

**Supplementary Figure 7.**
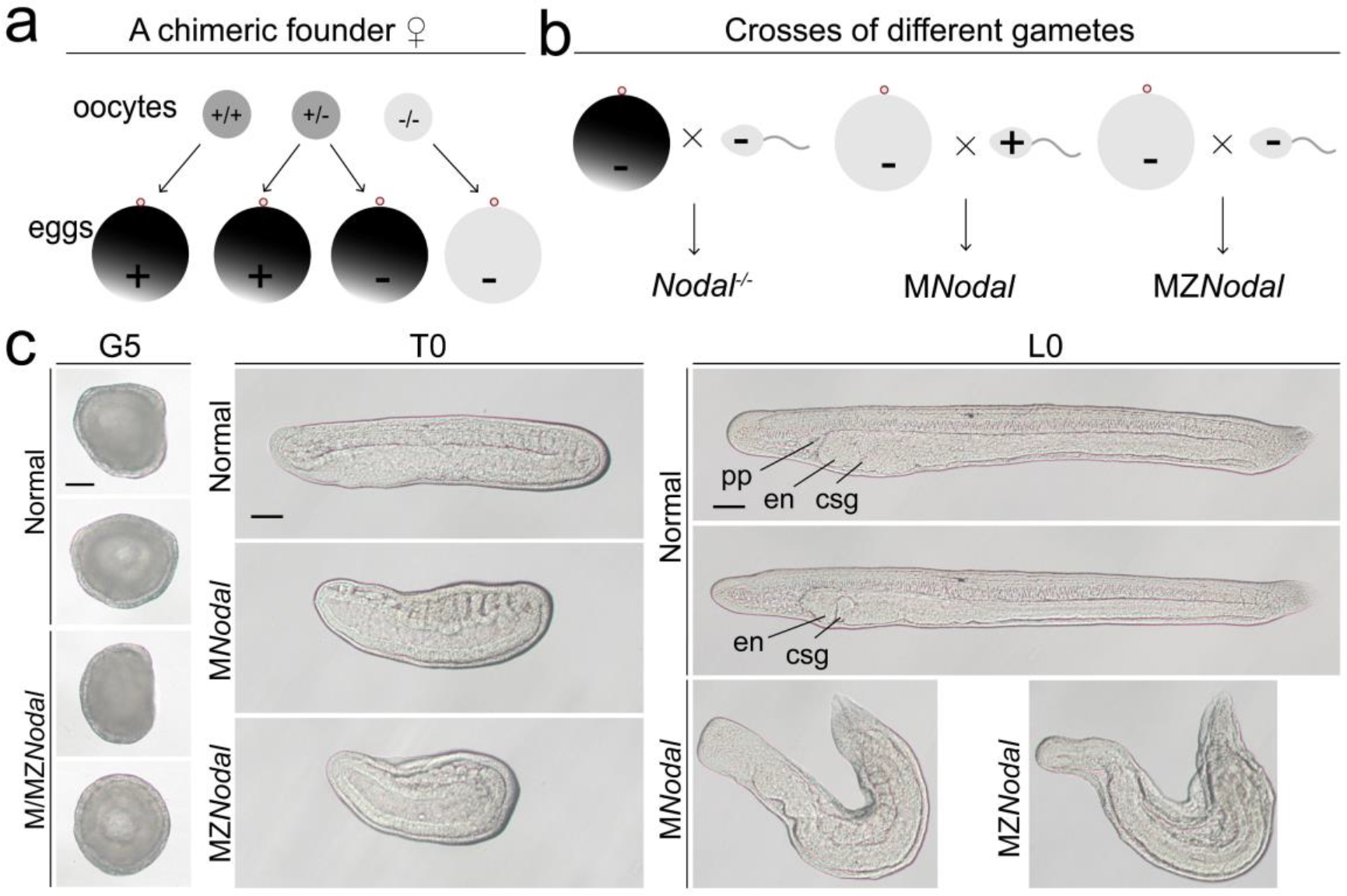
Phenotypic analysis of maternal and maternal-zygotic *Nodal* mutants. **(a)** Schematic diagram shows how the eggs without maternal *Nodal* accumulation are acquired from chimeric female founders injected with TALEN or Cas9/sgRNA. Dark grey indicates deposited maternal *Nodal* mRNA, while light grey marks no maternal *Nodal* mRNA accumulation. **(b)** Schematic diagram shows how zygotic (*Nodal^-/-^*), maternal (M*Nodal*) and maternal-zygotic *Nodal* (MZ*Nodal*) mutants used in the study are generated. **(c)** Phenotypic analysis of maternal and maternal-zygotic *Nodal* mutants. Embryos at G5 stage were viewed from the left side (upper panels) or blastopore (lower panels), with dorsal to the top. Embryos at other stages were viewed from the left side with anterior to the left. pp, preoral pit; en, emdostyle; csg, club shaped gland. Scale bars, 50 μm.

**Supplementary Figure 8.**
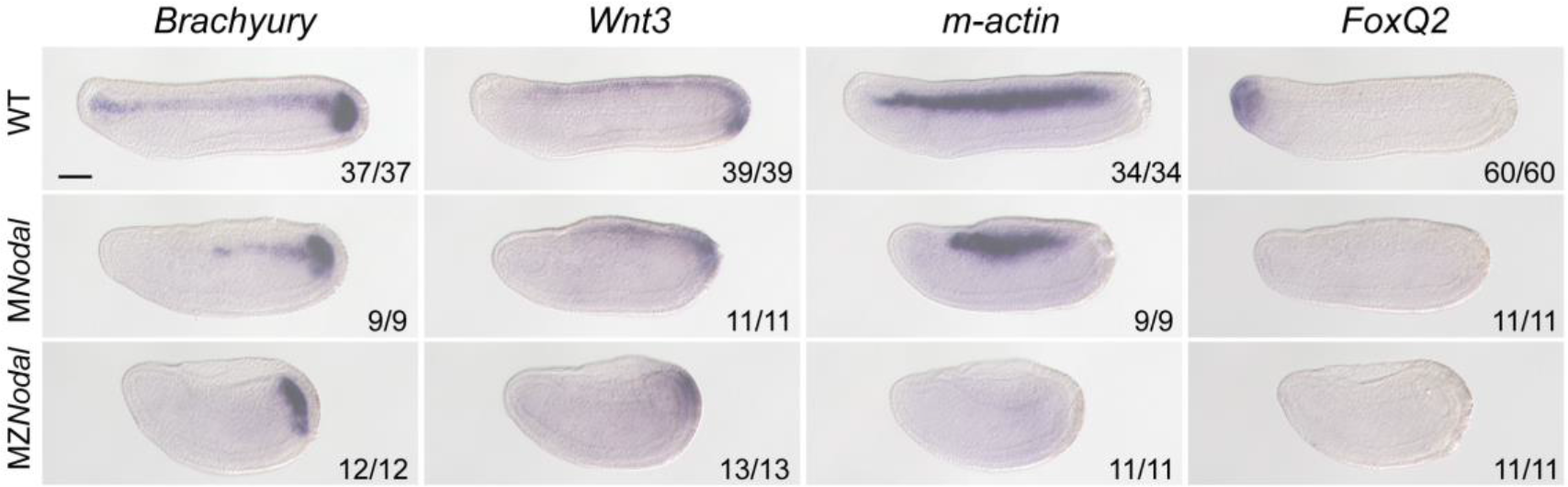
The expression of marker genes in maternal and maternal-zygotic *Nodal* mutants at T0 stage. All embryos were viewed from the left side with anterior to the left. M*Nodal* and MZ*Nodal* mutants were distinguished from their siblings based on their phenotypes. Numbers at the bottom right of each panel indicate the number of times the expression pattern shown was observed, out of the total number of examined embryos. Scale bar, 50 μm.

**Supplementary Figure 9.**
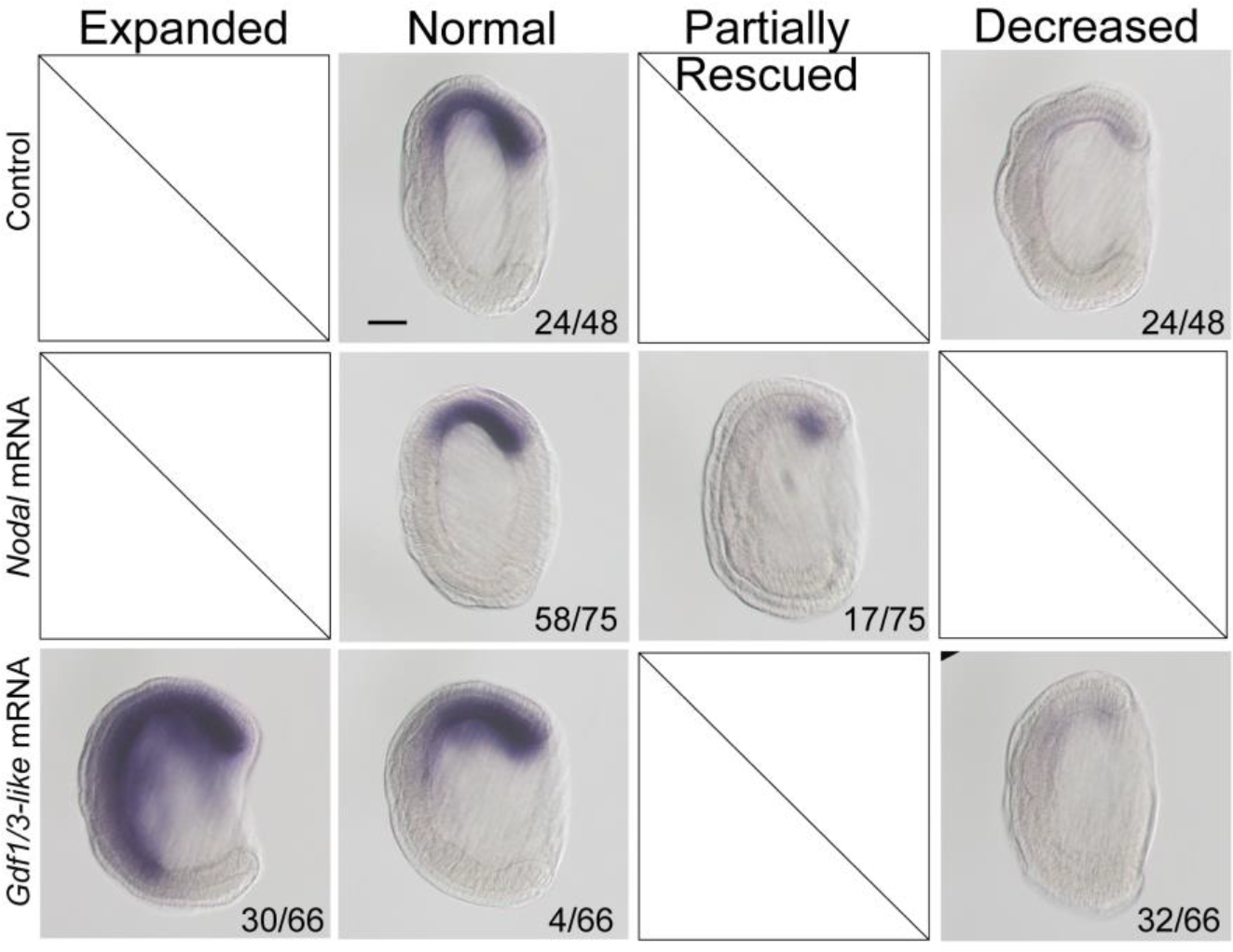
The expression of *Gsc* in maternal *Nodal* mutants injected with *Nodal* or *Gdf1/3-like* mRNA. Eggs from the female founder 1 (F0♀1) were respectively injected with *Nodal* and *Gdf1/3-like* mRNA, and then fertilized with sperms from a WT male. Uninjected eggs from the same animal were also fertilized with sperms from the same WT male and used as control. They were collected at G5 stage for *in situ* analysis. In control embryos, two types of *Gsc* expression pattern (normal and decreased) were observed, while in *Nodal* mRNA and *Gdf1/3-like* mRNA injected embryos, two (normal and partially rescued) and three (expanded, normal and decreased) types of *Gsc* expression pattern were respectively found. The number of embryos showing each *Gsc* expression pattern out of the total number of examined embryos are shown in the bottom of each panels. All embryos were viewed from the left side with anterior to the left. Scale bar, 50 μm.

**Supplementary Figure 10.**
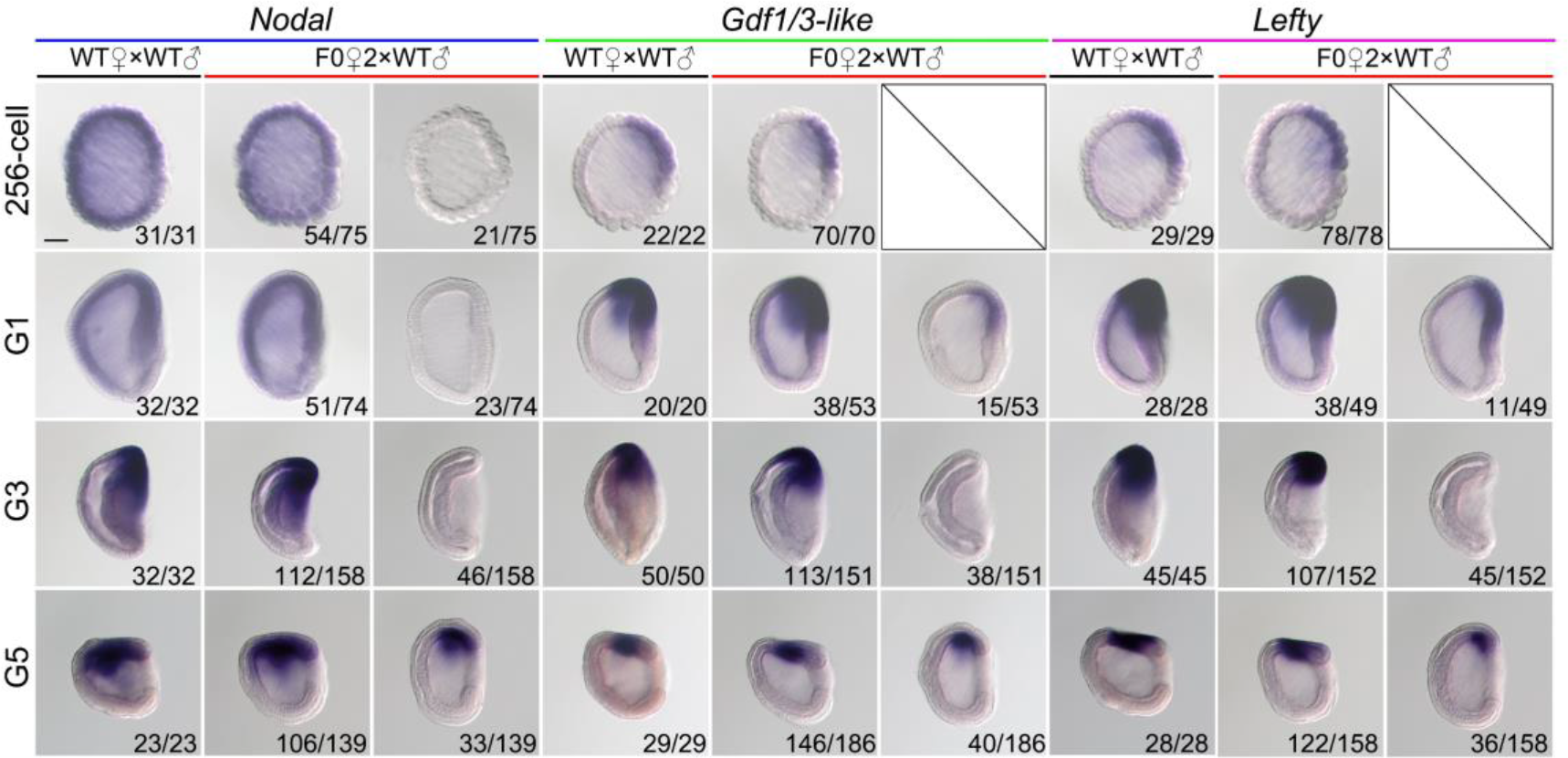
The expression of *Nodal*, *Gdf1/3-like* and *Lefty* in maternal *Nodal* mutants at four different stages. All embryos were viewed from the left side with anterior to the left. Numbers at the bottom right of each panel indicate the number of times the expression pattern shown was observed, out of the total number of examined embryos from each cross. Scale bar, 50 μm.

**Supplementary Table 1.**
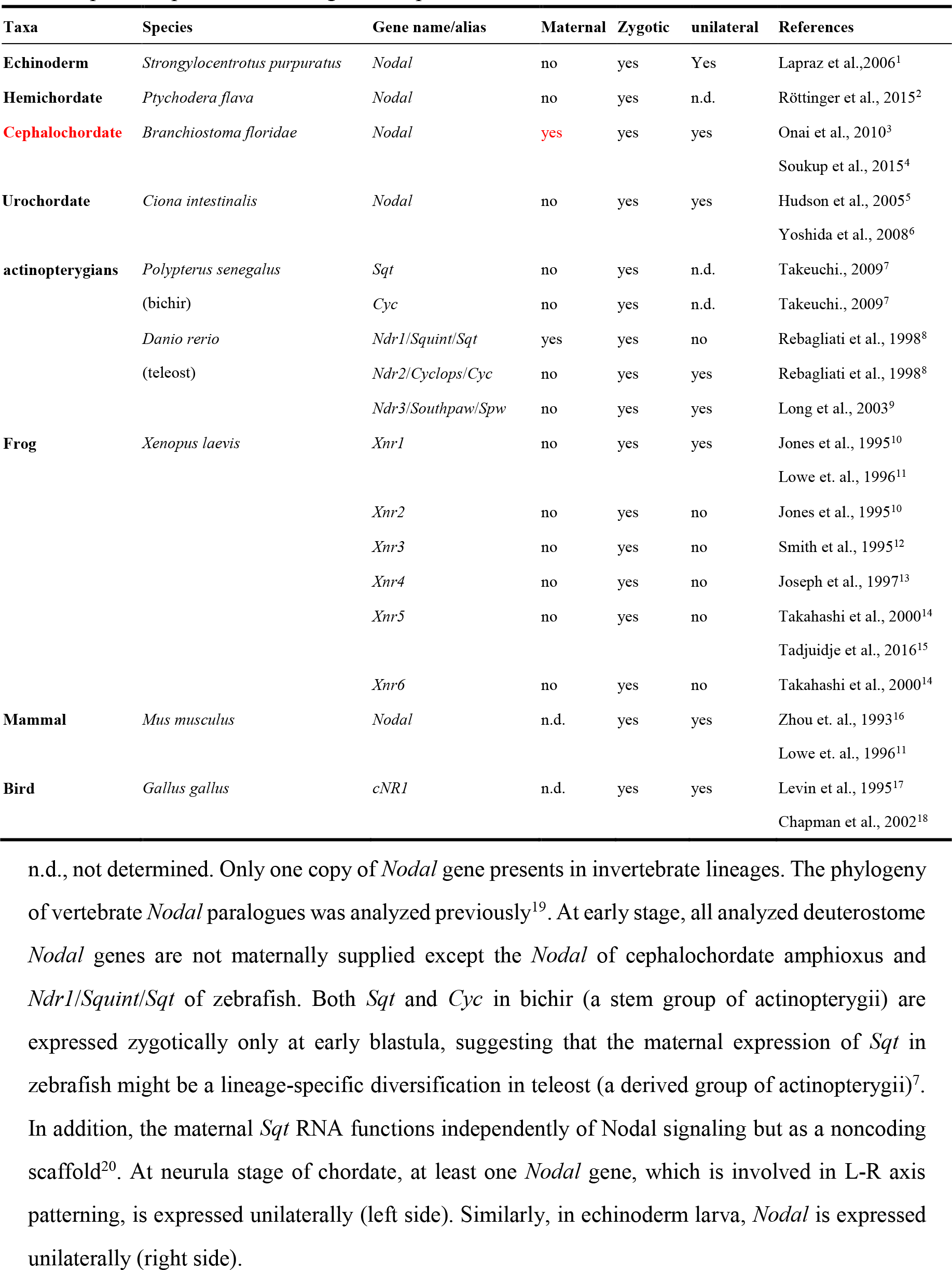
The expression pattern of *Nodal* gene in representative deuterostomes.

**Supplementary Table 2.**
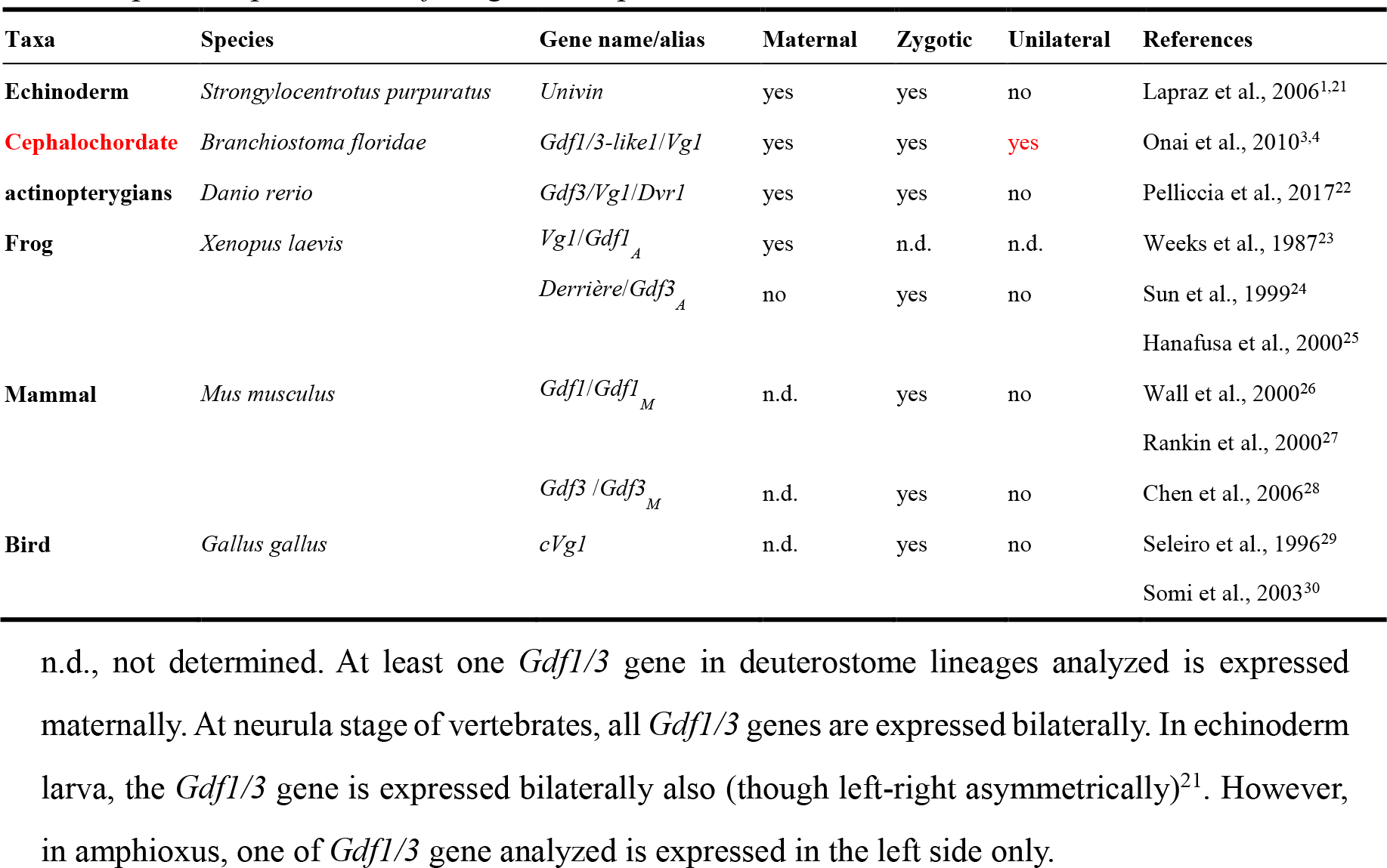
The expression pattern of *Gdf1/3* gene in representative deuterostomes.

## Notes

### Competing Interest Statement

The authors have declared no competing interest.

